# Independent modes of ganglion cell translocation ensure correct lamination of the zebrafish retina

**DOI:** 10.1101/066761

**Authors:** Jaroslav Icha, Christiane Grunert, Mauricio Rocha-Martins, Caren Norden

**Affiliations:** Max Planck Institute of Molecular Cell Biology and Genetics, Pfotenhauerstraße 108, 01307 Dresden, Germany; Instituto de Biofísica Carlos Chagas Filho, Av. Carlos Chagas Filho 373, 21941-902 Rio de Janeiro, Brazil

**Author notes:** Corresponding Author: Caren Norden, MPI-CBG, Pfotenhauerstraße 108, 01307 Dresden, Germany.

## Abstract

The arrangement of neurons into distinct layers is critical for neuronal connectivity and function of the nervous system. During development, most neurons move from their birthplace to the appropriate layer, where they polarize. However, kinetics and modes of many neuronal translocation events still await exploration. Here, we investigate ganglion cell (RGC) translocation across the embryonic zebrafish retina. After completing their translocation, RGCs establish the most basal retinal layer where they form the optic nerve. Using *in toto* light sheet microscopy, we show that somal translocation of RGCs is a fast and directed event. It depends on basal process attachment and stabilized microtubules. Interestingly, interference with somal translocation induces a switch to multipolar migration. This multipolar mode is less efficient but still leads to successful RGC layer formation. When both modes are inhibited, RGCs that fail to translocate induce lamination defects, indicating that correct RGC translocation is crucial for subsequent retinal lamination.

## Introduction

A conserved feature of the central nervous system (CNS) is its stratified organization. The accurate localization of neuronal subtypes into layers is established during development and is critical for the timely formation of connections among particular neurons. It thereby helps to ensure CNS functionality. Since many neurons are born in defined proliferative zones, from which they need to reach their final location, neuronal lamination relies on correct neuronal translocation. Neuronal movements are dynamic and often depend on the surrounding environment (Cooper, 2013; Marin et al., 2010). Therefore, they are best understood by time lapse *in toto* imaging experiments (Driscoll and Danuser, 2015). Despite this, insights about neuronal migration have often been generated using fixed tissue or *ex vivo* culture (Cooper, 2013). This is because many established model systems for studying neuronal migration, e.g. cerebellar granule neurons of rodents, are not easily imaged in intact embryos. Even though *in toto* imaging of neuronal translocation has been achieved in mouse embryos (Yanagida et al., 2012), the applicability of this experimental setup is limited. Consequently, model systems that allow live imaging in intact developing embryos need to be explored and findings there can then be used to understand neuronal translocation events in less accessible parts of the CNS.

The zebrafish (*Danio rerio*) is an ideal model system for live imaging approaches due to its small size and translucency at embryonic stages. Part of its CNS, the retina, is located at the surface of the animal, which makes it suitable for studying neuronal translocation and lamination. The retina consists of stereotypical layers of five well-defined neuronal cell types (photoreceptors, horizontal, bipolar, amacrine and ganglion cells) and one glial cell type (Cajal, 1972; Galli-Resta et al., 2008). These layers arise between 30 and 72 hours post fertilization (hpf), which means that a single imaging experiment can cover the complete time frame of neurogenesis. However, it is important to choose the right microscope depending on the question asked. Whereas confocal microscopy can induce phototoxicity-related artifacts during zebrafish development (Jemielita et al., 2012) and is usually performed at lower temporal resolution (Chow et al., 2015; Poggi et al., 2005), light sheet fluorescence microscopy (LSFM) allows fast, less phototoxic imaging (Huisken et al., 2004; Stelzer, 2015). Consequently, LSFM is an ideal tool to image the emergence of neuronal layering in the zebrafish retina.

The first neuronal lamination event in the retina is the formation of the retinal ganglion cell (RGC) layer. RGCs are the first retinal neurons born in vertebrates (Cajal, 1972; Nawrocki, 1985; Sidman, 1961). Upon their apical birth, RGCs move basally, spanning the complete apico-basal axis towards the lens, where their axons later form the optic nerve. Although it is known that RGCs can translocate by moving their soma while keeping the attachment to both apical and basal sides of the epithelium (Cajal, 1972; Hinds and Hinds, 1974; Zolessi et al., 2006), we still lack a thorough analysis of their translocation kinetics and modes. Additionally, the cell biological mechanisms of somal translocation are still unknown, despite the fact that this is a widespread mode of neuronal movement during retinal (Chow et al., 2015) and brain development (Nadarajah et al., 2001).

Based on high-resolution imaging of single cell behavior, we analyze the kinetics of RGC translocation. We show that RGC somal translocation is an active, directionally persistent process. Speed and directional persistence of somal translocation depend on intact microtubules (MTs) and basal cellular attachment. Interestingly, interference with somal translocation can result in a switch to a multipolar migratory mode, not previously reported for RGCs. The multipolar mode is less efficient, but allows RGCs to reach their correct location. Complete inhibition of RGC movements, however, substantially perturbs lamination of other retinal layers. Thus, our study sheds new light on the mechanisms of and requirements for efficient basal translocation of RGCs and highlights its importance for subsequent retinal lamination.

## Results

### Rapid RGC translocation is followed by a period of fine positioning

To study the emergence of the RGC layer as well as the kinetics of RGC translocation, we labeled these cells using constructs containing the *ath5 (atonal bHLH transcription factor 7, atoh7*) promoter (Fig. 1A) (Brown et al., 1998; Masai et al., 2003). Ath5 starts to be expressed during the last cell cycle in a subset of progenitor cells. These progenitors divide apically and produce one RGC and one cell that later gives rise to photoreceptors (Poggi et al., 2005; He et al., 2012). To label and follow RGCs between their apical birth and axonogenesis, we made use of previously published *Tg(ath5:GAP-GFP/RFP)* lines (Zolessi et al., 2006), which express membrane-targeted GFP or RFP in the *ath5* lineage (Fig. 1 A and B). Alternatively, to achieve mosaic labeling and follow single RGCs, we injected *ath5* plasmid DNA into one cell stage embryos, typically *ath5:GFP-CAAX* coding for membrane targeted GFP (for example Fig. S1, G–I). We noted that the use of spinning disk microscopy even at low intensity illumination induced a slowing down of RGC translocation compared to a LSFM setup (Fig. 1 C and S1 A). The two microscope setups differ in the way the sample is mounted and in the amount of light the sample is exposed to during image acquisition. To test the influence of mounting, we compared the viability and development of embryos mounted for LSFM or spinning disk microscopy to non-mounted embryos (Fig. S1 B). We observed no change in heart rate or apoptosis (Fig. S1, C–E) between the samples, therefore we concluded that the observed differences in RGC translocation speed at different microscopes are likely due to higher phototoxicity induced by spinning disk microscopy. For this reason, we used only LSFM-generated data to analyze RGC translocation kinetics. We confirmed that this technique does not produce noticeable phototoxicity-related artifacts by comparing embryos imaged for 12–16 hours in the LSFM to non-imaged controls. In both conditions the RGC layer formed to a similar extent.

**Figure 1.**
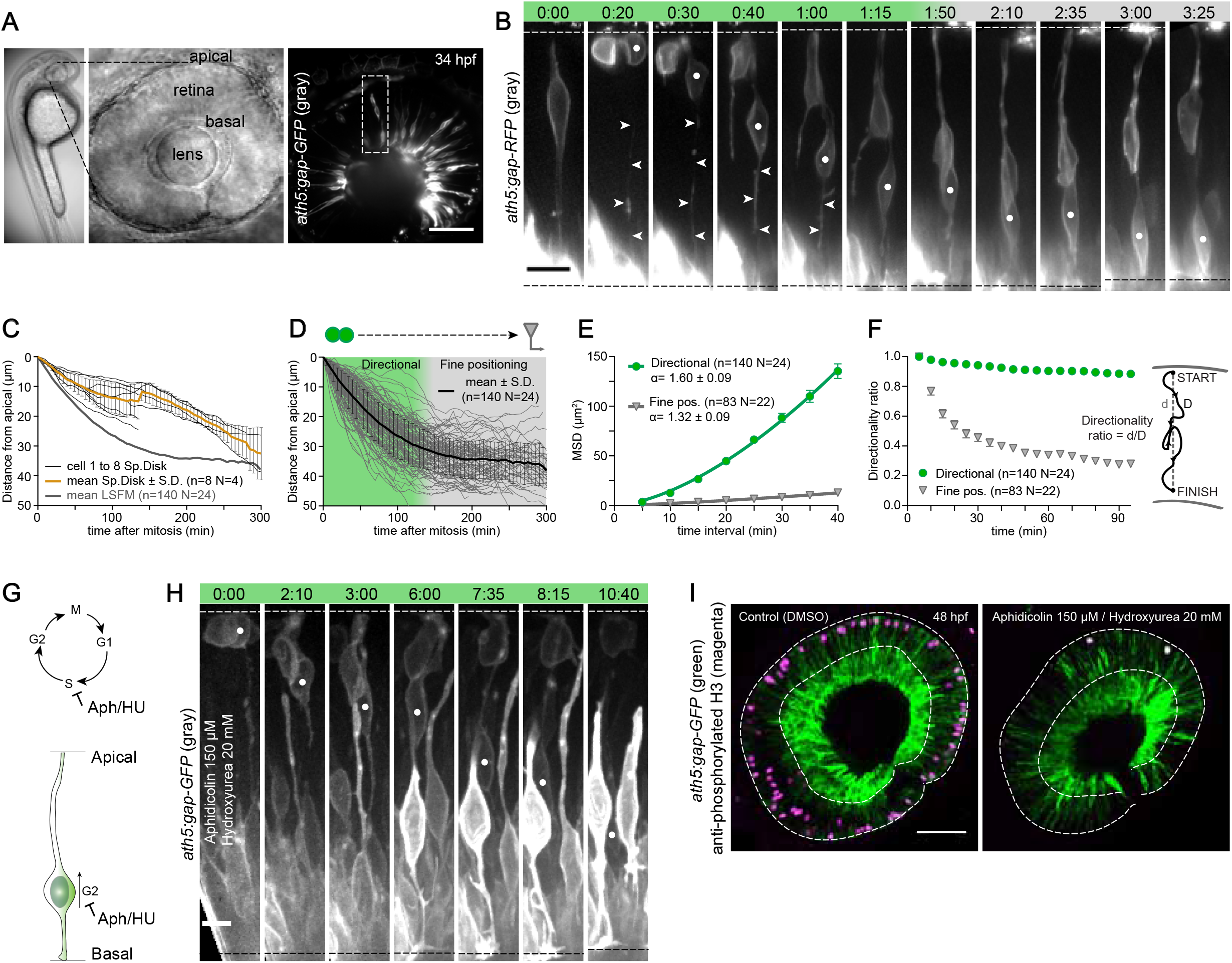
RGC translocation kinetics; RGC somal translocation can be divided into directional and fine positioning phases and is not dependent on cell cycle of neighboring progenitors. A) Developing eye of a 34 hours post fertilization (hpf) embryo. *ath5:gap-GFP* transgene labels RGCs. The dashed box shows the typical area displayed in subsequent montages. In all figures the apical side of the retina is up and the basal side is down. Scale bar: 50 μm. B) Typical example of RGC translocation in LSFM. Green phase: directionally persistent movement. Gray phase: fine positioning. Dashed lines delimit the apical and basal sides of the retina. White dot: RGC followed. Arrowheads: basal process. Time is shown in hh:mm. Scale bar: 10 μm. C) Kinetics of RGC translocation in a spinning disk confocal microscope. 0 indicates mitotic position of cells. All trajectories in spinning disk microscope (n=8 cells, N=4 experiments) and an average trajectory ±S.D. are shown plus the average of WT trajectories in LSFM. The jump in the average value at 140 min is caused by averaging fewer values. D) Kinetics of RGC translocation in LSFM. 0 indicates mitotic position of cells. 300 min range represents the average time from mitosis to axonogenesis. 140 single trajectories and an average trajectory ±S.D. are shown. Green phase: directionally persistent movement. Gray phase: fine positioning. E) MSDs of RGCs in directional and fine positioning phases. Directional phase: n=140 N=24. Fine positioning: n=83 N=22. MSDs were calculated from the first 95 min after mitosis and the first 95 min after reaching the basal side. The α-value is given with 95% confidence interval. Error bars represent S.E.M. F) Directionality ratio of RGCs in directional and fine positioning phases. The same data as in Fig. 1 E are used. The average of all tracks is shown with error bars representing S.E.M. Directionality ratio at the end of the trajectory: Directional=0.88 Fine pos.=0.28. The scheme: directionality ratio is defined as a ratio between the distance from start to finish of the trajectory (d) and the length of the actual trajectory (D). G) Aphidicolin/Hydroxyurea stalls cells in S phase. Thus, progenitors do not enter G2, and their nuclei do not migrate towards the apical side for mitosis. H) RGC translocation still occurs after cell cycle inhibition. An *ath5:gap-GFP* transgenic embryo was imaged by spinning disk microscopy with 150 μM Aphidicolin/20 mM Hydroxyurea added around 34 hpf. Imaging was started 1 hour after drug addition. Time is shown in hh:mm. Dashed lines delimit the apical and basal sides of the retina. Scale bar: 5 μm. I) The RGC layer still forms after cell cycle inhibition. Fewer mitotic cells (right) compared to control (left) were observed by pH3 staining (magenta). Dashed lines mark retinal outline and RGC layer. Scale bar: 50 μm.

Following emerging RGCs by live imaging revealed that RGC translocation was divided into two parts that in total took around 285 min (median, n=50 cells N=19 experiments) (Fig. 1 B and D). Initially, cells translocated basally for 115 min (median, n=140 N=24) in a fast and persistent manner, moving about 28 μm away from the apical side (median, n=140 N=24) (Fig. 1 B and D). After this phase, RGCs exhibited slower, more random movements within the RGC layer in the basal part of the retina for 165 min (median, n=50 N=19) (Fig. 1 D and S1 F) moving on average additional 3 μm basally. This probably reflects the growth of the retina rather than net cell movement. We termed this second phase fine positioning. It ended with axonogenesis, which we defined as the moment when a persistent growth cone emerged from the RGC. Only during fine positioning, cells lost their apical process, around 60 min before axonogenesis (median, n=33 N=16), although the exact timing was variable (Fig. S1 F).

To quantify the kinetics and directional persistence of the two phases of RGC movement, we calculated mean squared displacements (MSD) (Leung et al., 2011); (Okamoto et al., 2014) from manually tracked 2D trajectories (Fig. 1 E). MSD analysis allows determining the directionality of a particle (here the nucleus) from the shape of the MSD curve. A linear curve indicates random motion and a supralinear curve indicates directional motion of the particle. Our MSD analysis showed that the initial basal movement of RGCs was directionally persistent as indicated by the supralinear MSD and the α value of 1.60±0.09 from fitting the MSDs with 2D_Δ_t^α^ (see Materials and Methods). In contrast, MSDs of cells undergoing fine positioning showed an almost linear relationship with time with an α value of 1.32±0.09 (Fig. 1 E), indicating random motion. We also calculated the directionality ratio of the trajectories. Here, values close to 1 indicate high directionality and values approaching 0 indicate random motion (Fig. 1 F). This analysis confirmed that initial basal translocation is much more directed than fine positioning (Fig. 1 F). RGCs in the directional phase moved with a median instantaneous velocity of 0.26 μm/min (Fig. S1 A). This was calculated from 1D apico-basal displacement, where a positive sign was assigned to basalward movement and a negative sign to apicalward movement. Median instantaneous velocity of fine positioning was close to 0 (0.02 μm/min), as expected for a cell undergoing random motion (Fig. S1 A). We conclude that RGC translocation to the basal side of the retina before RGC layer formation consists of a directional and a fine positioning phase.

### Basal movement of RGC and progenitor nuclei is more efficient when cells inherit the basal process

So far, our analysis showed that initial RGC translocation is directionally persistent. Two scenarios could explain how this directionality arises: a) RGCs are displaced basally by the surrounding progenitor nuclei undergoing directed apical migration, as suggested for the gradual basal displacement of progenitor cell nuclei in the proliferative phase (Norden et al., 2009); or b) RGC movements are driven cell autonomously and occur independently of the movements within surrounding cells.

To test option a), we stalled cells in S-phase using a combination of Aphidicolin and Hydroxyurea (Fig. 1 G). We started cell cycle inhibition at 32 hpf, when the first pool of RGCs has been born and co-inhabits the retina together with an abundant pool of progenitor cells (Kay, 2005). Apical migration of nuclei in these progenitor cells relies on progression through the cell cycle and has been shown to be stalled in this condition (Leung et al., 2011). RGCs were either analyzed live (Fig. 1 H) or fixed at 48 hpf, when the RGC layer was fully formed in control embryos (Fig. 1 I). Inhibition of proliferation was confirmed by the lack of anti-phosphorylated histone H3 staining in drug-treated embryos (Fig. 1 I). We noted that migration of RGCs still occurred (Fig. 1 H) and although the eyes of treated embryos were overall smaller, the RGC layer still formed (Fig. 1 I). This indicated that RGC translocation depended on cell intrinsic processes. Various active modes of neuronal displacement are driven by the concerted action of the centrosome and associated Golgi apparatus. For example, it was shown that during glial guided neuronal migration the centrosome leads the translocation process and pulls the nucleus via a MT cage (Solecki et al., 2004; Bellion et al., 2005; Tsai et al., 2007). However, it was previously suggested that in RGCs the centrosome remains in the apical process during translocation (Zolessi et al., 2006; Hinds and Hinds, 1974). To analyze how centrosome and Golgi behave during RGC displacement, we used Centrin-tdTomato as a centrosomal marker and the trans-Golgi marker GalT-RFP. Live imaging showed that both structures remain in the apical process during somal translocation and only follow the nucleus towards more basal positions once the apical process is retracted during the fine positioning phase in all cells followed (Fig. S1 G and S1 I). Similar observations were made for the primary cilium marked by Arl13b-mKate2 in all cells followed (Fig. S1 H) (Lepanto et al., 2016). This suggests that these organelles do not actively lead nucleokinesis and somal translocation in RGCs.

Given that RGC displacement was not triggered by movement within surrounding cells and was not driven by a centrosome based mechanism, we turned our attention to the basal process as a guide for RGC translocation. In the mouse cerebral cortex, the inheritance of the basal process has been shown to streamline basal displacement of progenitor cell nuclei (Okamoto et al., 2013). We thus investigated how basal process inheritance correlates with RGC displacement kinetics. Interestingly, the basal process was inherited preferentially by the RGC in 78% of divisions (n=109/140). In the remaining 22% RGCs regrew the basal process later during their translocation. Translocation kinetics of RGCs inheriting the basal process was faster than for RGCs that had to regrow it (compare Fig. 1 B and 2 A). As seen in representative trajectories (Fig. 2 B; more trajectories in Fig. S2, F and G), RGCs not inheriting the basal process (n=31/140 cells) moved basally more slowly (Fig. 2, E and F). The higher directional persistence of RGCs inheriting the basal process was apparent from MSDs (Fig. 2 G and Fig. S2 A), directionality ratios (Fig. 2 J) and instantaneous velocity distributions (Fig. S2 C). Even though the cells not inheriting the basal process took longer to reach basal positions (165 vs. 105 min, median) (Fig. S2 E), the time interval between their apical birth and axonogenesis was comparable to the timing in cells inheriting the process (Fig. S2, D and E).

**Figure 2.**
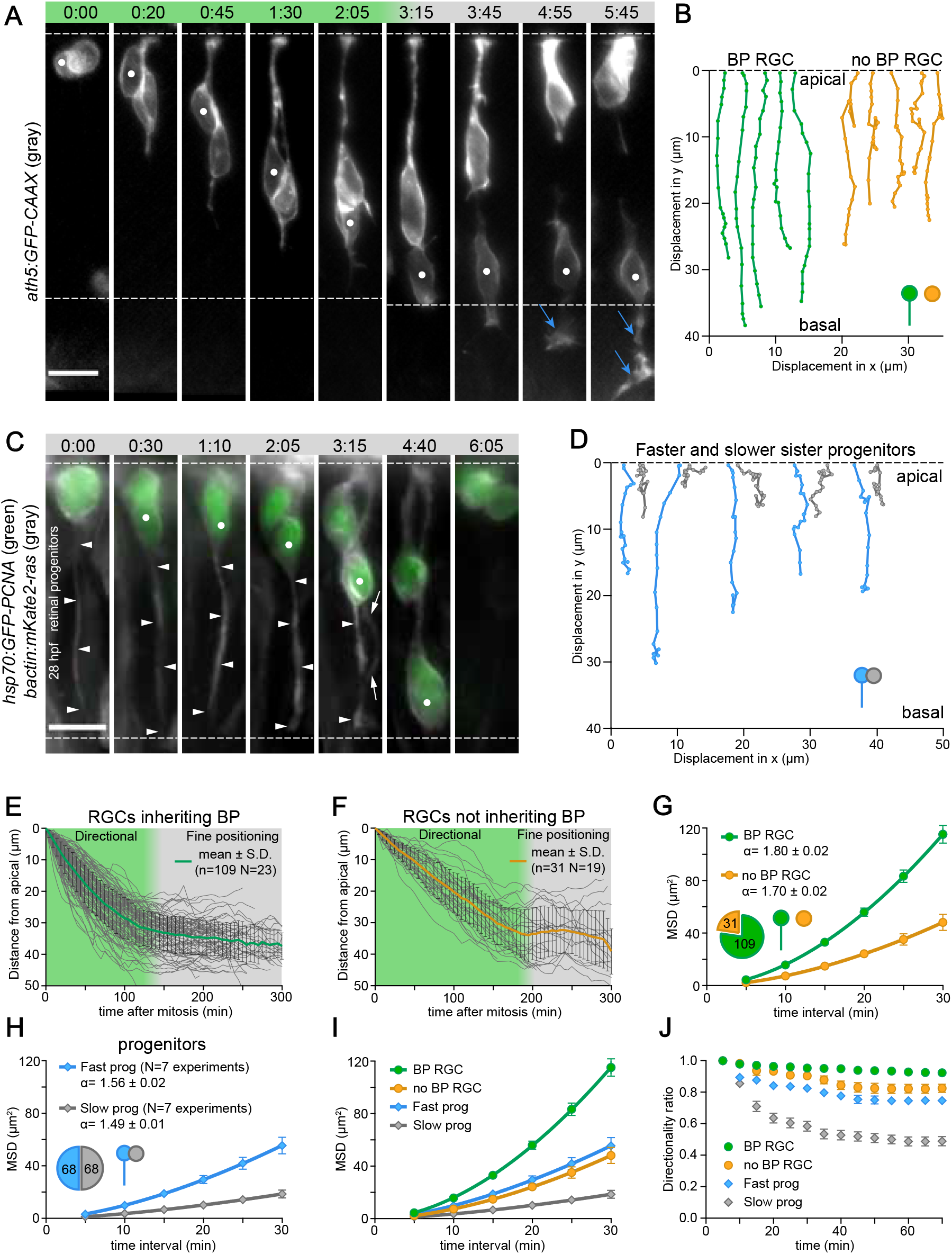
Basal process inheritance streamlines basal nuclear translocation in RGCs and progenitors. A) Translocation of an RGC not inheriting the basal process (BP): compare to Fig. 1 B. Green phase: directionally persistent movement. Gray phase: fine positioning. Dashed lines delimit the apical and basal sides of the retina. White dot: RGC followed. Blue arrow: axon. Time is shown in hh:mm. Scale bar: 10 μm. B) Five representative 2D trajectories of RGCs inheriting (green) and RGCs not inheriting the basal process (yellow) for the first 95 min after cell division. For more trajectories see Fig. S2, F and G. C) Inheritance of basal process in progenitors. Dashed lines delimit the apical and basal sides of the retina. White dot: basal process inheriting progenitor. Arrowheads: inherited BP. Arrows: newly formed BP of the sister cell. Scale bar: 10 μm. D) Five representative 2D trajectories of sister progenitors inheriting (blue) and not inheriting the basal process (gray) for the first 95 min after cell division. E) Kinetics of RGC translocation with basal process. 0 indicates mitotic position of cells. Single trajectories and an average trajectory ±S.D. are shown. Green phase: directionally persistent movement. Gray phase: Fine positioning. F) Kinetics of RGC translocation without basal process. G) MSDs of translocating RGCs with and without basal process. The MSDs were calculated from the first 70 min after mitosis. The cells with and without the basal process are independent (not sister cells). The α-value is given with 95% confidence interval. Error bars represent S.E.M. H) MSDs of translocating faster and slower sister progenitor nuclei. The MSDs were calculated from the first 70 min after mitosis. The α-value is given with 95% confidence interval. Error bars represent S.E.M. I) A comparison of MSDs of RGC and progenitor nuclear translocation. The graph shows combined data from (G) and (H). J) Directionality ratios of RGC and progenitor nuclear translocation. The average of all tracks is shown with error bars representing S.E.M. Directionality ratio at the end BP RGC= 0.92, no BP RGC= 0.82, Fast prog= 0.75, Slow prog= 0.49.

Retinal progenitors share many morphological similarities with RGCs. Thus, we hypothesized that a positive effect of basal process inheritance on basal nuclear displacement might also exist after apical progenitor divisions. While it was previously reported that basal displacement of retinal progenitor nuclei fits the characteristics of a random walk (Leung et al., 2011; Norden et al., 2009), these analyses did not discriminate between progenitors with or without a basal process. To test whether progenitors show different kinetics of basal nuclear translocation depending on whether they inherit the basal process, we analyzed them by labeling with the nuclear marker PCNA-GFP and mKate2-ras to mark membranes (Fig. 2 C). Velocities by which nuclei of sister cells moved away from the apical side differed in all instances followed (Fig. S2 C). The faster nucleus of the pair also consistently showed higher directional persistence and directionality ratio (Fig. 2, H-J and Fig. S2 B). These nuclei were followed for the first 70 minutes after cell division, before they entered a more stochastic movement mode (Leung et al., 2011). The trajectories of the sister nuclei (n=68 pairs N=7) moving away from the apical side differed in all instances followed (see examples in Fig. 2 D). One of the nuclei consistently showed higher directional persistence (Fig. 2 H and Fig. S2 B), directionality ratio (Fig. 2 J) and instantaneous velocity distributions (Fig. S2 C). In almost all (16/17) cases in which we could confidently assign basal process inheritance to one of the progenitors, the faster and more directionally persistent nucleus belonged to the progenitor cell that inherited the basal process. Nevertheless, in comparison to RGCs the movement of nuclei in progenitors was less directional and slower (Fig. 2, I and J and Fig. S2 A–C). Overall, these data indicate that basal process inheritance is an important factor for efficient basal nuclear translocation in RGCs as well as in progenitors. The fact that nuclear displacement in progenitors is slower than RGC displacement suggests that RGC translocation involves additional components.

### Interference with basal process attachment impairs basal RGC movement

As basal process inheritance streamlined basal RGC movement, we tested how interference with basal process attachment to the basal lamina affects RGC translocation. To achieve this, we initially interfered with the basal lamina itself. The basal lamina consists of extracellular matrix (ECM) proteins with Laminin α1 being a core component (Fig. S3 A) (Randlett et al., 2011b). The knock down of *laminin α1* using a published morpholino (Randlett et al., 2011b) depleted Laminin from the basal retinal membrane (Fig. S3 B) and RGCs in this condition lost their basal process, which led to defective translocation and little net movement towards the basal side (Fig. S3 C). However, laminin depletion can have diverse effects on late retinal formation (Fig. S3 B) (Randlett et al., 2011b). Therefore, we additionally interfered with basal RGC attachment by targeting the basal actin pools, which are associated with focal adhesion and ensure cellular attachment to the ECM (Martinez-Morales et al., 2009, Randlett et al., 2011a). We used the Arp2/3 complex inhibitor CK-666 (Nolen et al., 2009) and the Rho kinase (ROCK) inhibitor Rockout. Arp2/3 inhibition has recently been shown to disrupt basal attachment of radial glia cells in the mouse cerebral cortex (Wang et al., 2016). We determined the necessary inhibitor concentrations at which the basal actin was reduced while actin associated with apical adherens junctions was still present (Fig. S3, D and E). In both treatments this led to basal process detachment (Fig. 3 A and 5F). In these conditions, RGCs did not stably reattach their basal process after their final apical division and did not persistently move basally (Fig. 3 A and 5F). In contrast, apical staining in corneal cells, where analysis is easier due to their bigger apical surface, Rockout treatment did not show major effects on cortical actin and microridges (Lam et al., 2015) while CK-666 treatment led to a disarray of microridges and apical area reduction (Fig. S3 F).

**Figure 3.**
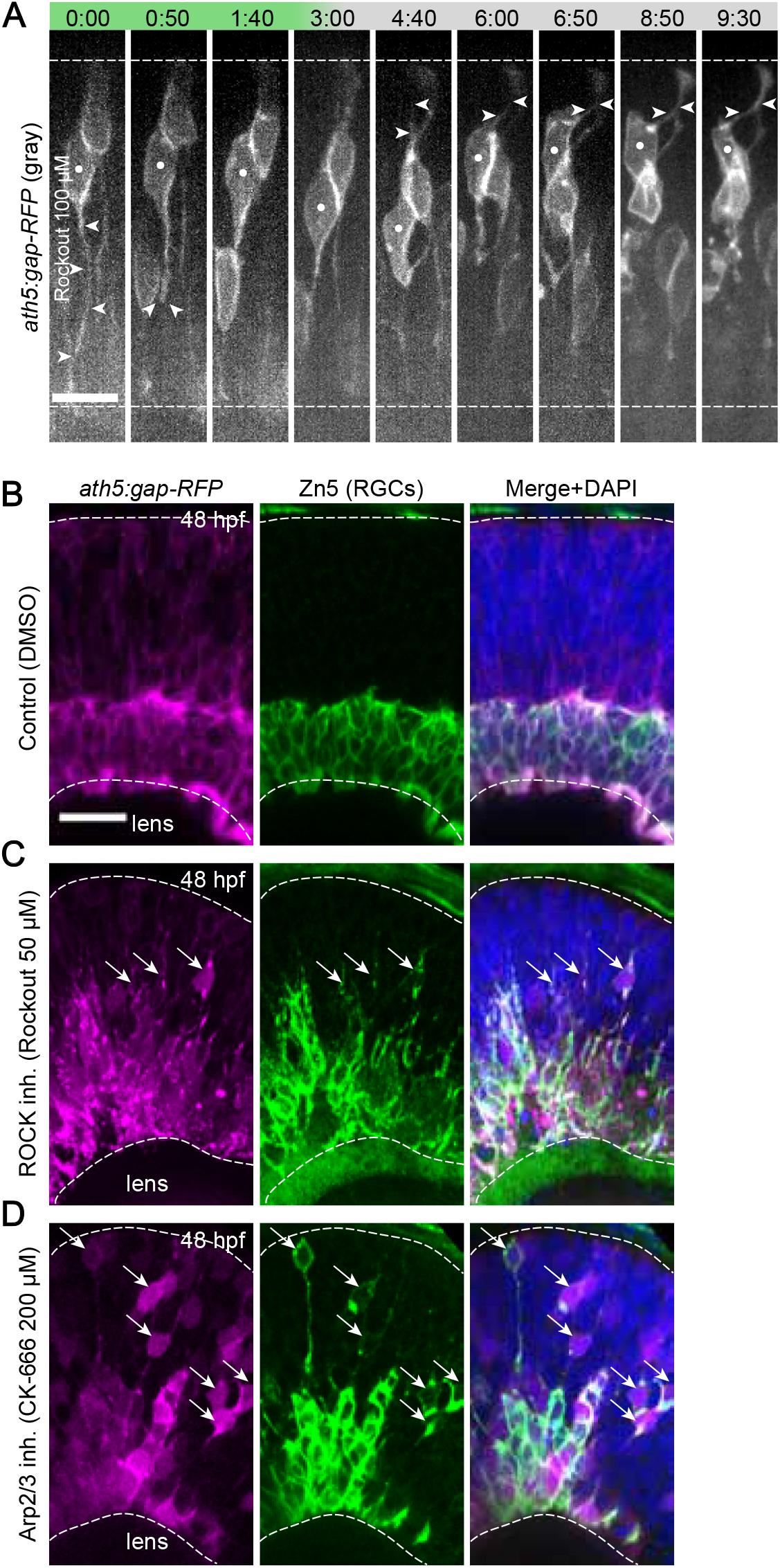
Basal process attachment is important for RGC translocation. A) Live imaging of RGC translocation after ROCK inhibition. The *ath5:gap-RFP* fish were imaged at a spinning disk microscope from 34 hpf. Rockout was added at the start of imaging. Note that the RGC loses the basal process but keeps the apical process. White dot: RGC followed. Arrowheads: basal and apical process. Time is shown in hh:mm. Dashed lines delimit the apical and basal sides of the retina. Scale bar: 10 μm. B) Staining for differentiated RGCs with Zn5 antibody in control retina at 48 hpf. C) ROCK inhibition interferes with RGC translocation. Staining for differentiated RGCs with Zn5 antibody after ROCK inhibition. D) Arp2/3 inhibition interferes with RGC translocation. Staining for differentiated RGCs with Zn5 antibody after Arp2/3 inhibition. B), C), D) Dashed lines delimit the apical and basal sides of the retina. Arrows: ectopic RGCs. Scale bar: 20 μm.

To investigate the long-term effects of these inhibitors on RGC displacement, we stained RGCs with a Zn5 antibody (Trevarrow et al., 1990) at 48 hpf, 16 hours after drug addition. Zn5 antibody labels differentiated RGCs independent of their location. In both conditions, ectopic RGCs were observed in the middle of the retina while in control retinas Zn5-positive RGCs had formed a compact layer at the basal side (Fig. 3, B–D). This means that cells differentiated into RGCs in time, even when they did not reach the most basal retinal layer. To substantiate our findings in the drug conditions and exclude pleiotropic effects we additionally used a genetic approach. We overexpressed a C-terminal domain of Arp2/3 complex activator N-WASP (NWASP-CA). This truncated protein binds the Arp2/3 complex, but does not activate it and thus acts as dominant negative that inhibits Arp2/3 activity (Rohatgi et al., 1999) similarly to the CK-666 drug. Also this approach affected the actin cytoskeleton (Fig. S3 F) and led to RGC translocation defects (Fig. 5 G–J and S4 H and I, discussed below), confirming that indeed the specific disruption of basal process attachment leads to impaired RGC translocation.

### Efficient RGC movement depends on a stabilized microtubule cytoskeleton

As we observed that basal displacement is more efficient for RGCs than for progenitor cells (Fig. 2, H and I), we asked which additional components could play a role in nuclear translocation of RGCs. An attractive candidate was the MT cytoskeleton, which is involved in many modes of neuronal translocation (Umeshima et al., 2007; Solecki et al., 2004; Cooper, 2013). To test how MTs behave during RGC translocation, we first compared their growth dynamics in RGCs and progenitors by using the plus tip marker End-binding protein 3 (EB3) (Stepanova et al., 2003) (Fig. 4 A). EB3 follows the plus tips of MTs, which allows for measurement of MT growth rate. The moving EB3 dots are termed ‘comets’. In both, progenitors and RGCs, the vast majority of EB3 comets emanated from the apical centrosome and moved basally as shown previously (Norden et al., 2009, Tsai et al., 2007) (Fig. 4 A). Notably, MT growth was faster in progenitors (0.23 μm/s, vs. 0.13 μm/s, median) (Fig. 4 A), demonstrating that there are differences in the MT growth rates between these two cell types. The slower comet speed in the RGCs was indicative of a more stabilized, less dynamic MT cytoskeleton in these cells. To test this idea, we stained embryos with anti-acetylated tubulin antibody as a read-out for MT stabilization (LeDizet and Piperno, 1986). Additionally, all MTs were labeled with Doublecortin *Tg(bactin:GFP-DCX)* (Distel et al., 2010), a MT-associated protein. We found that apical processes of translocating RGCs were rich in acetylated tubulin (Fig. 4 C) unlike the apical processes of progenitor cells (Fig. 4 B), which is in line with the slower EB3 comets in RGCs. To understand the emergence of these stabilized MTs, we examined them live in RGCs using *ath5* promoter-driven Doublecortin *ath5:GFP-DCX*. MTs became enriched in the apical process shortly after cell division and remained there during the whole translocation process (Fig. 4 D), as suggested before analysing electron microscopy images (Hinds and Hinds, 1974).

**Figure 4.**
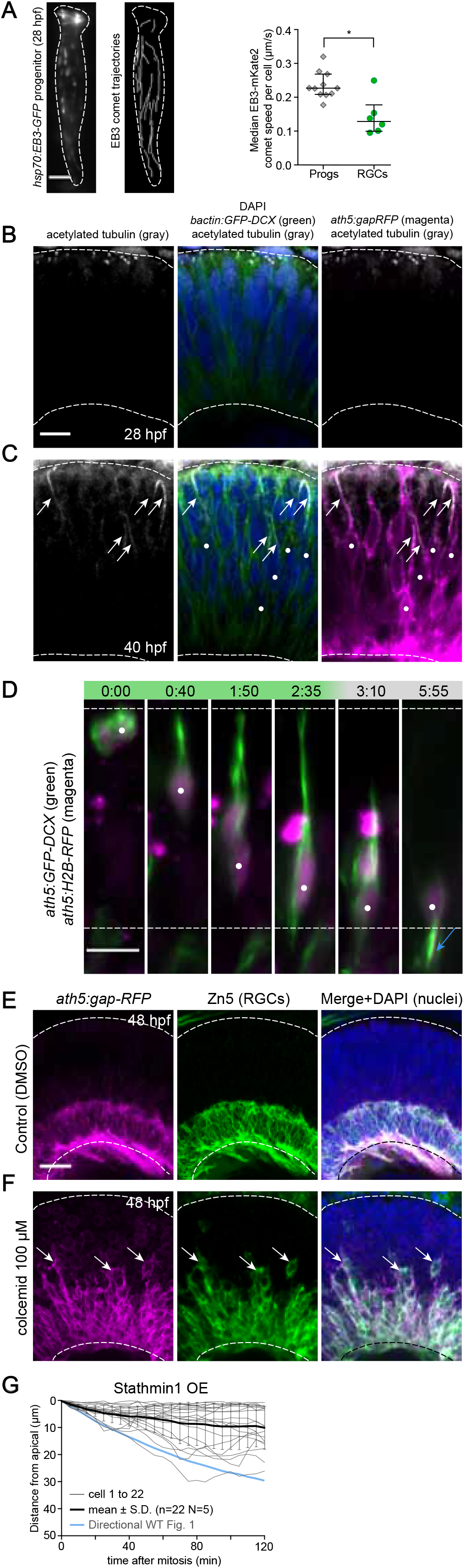
Stabilized microtubules are important for RGC translocation. A) EB3 comets are faster in progenitors than in RGCs. Left: an illustrative example of the EB3 data in a progenitor cell; scale bar: 5 μm. Right: progenitor speed median=0.23 μm/s n=11 cells, 60 comets. RGC speed median=0.13 μm/s n=6 cells, 24 comets, p=0.0103 Mann Whitney U test. Speed was measured 4 hours after heat shock in progenitors at 28 hpf, in RGCs at 38 hpf. The bars represent median and interquartile range. B) Acetylated tubulin antibody staining in progenitors. All microtubules were labeled by *bactin:GFP-DCX*. At 28 hpf only primary cilia at the apical side of progenitors are stained. Dashed lines represent apical and basal sides of the retina. Scale bar: 10 μm. C) Acetylated tubulin antibody staining in RGCs. All microtubules were labeled by *bactin:GFP-DCX*. At 40 hpf acetylated tubulin is seen in the AP of RGCs (white arrows). White dot: translocating RGCs. Dashed lines represent apical and basal sides of the retina. D) Live imaging of microtubules in translocating RGCs. Note microtubule accumulation in the apical process. Time is shown in hh:mm. Dashed lines delimit the apical and basal sides of the retina. White dot: RGC followed. Blue arrow: axon. Scale bar: 10 μm. E) Staining for differentiated RGCs with Zn5 antibody in control retina at 48 hpf. F) Zn5 staining in retinas treated with colcemid. E), F) Arrows: ectopic RGCs. Dashed lines delimit the apical and basal sides of the retina. Scale bar: 20 μm. G) Microtubule destabilization by overexpression of Stathmin1 *(hsp70:Stathmin1-mKate2)* stalls RGC translocation. Fish were heat shocked at 30 hpf and imaged from 34 hpf. Graph shows all trajectories after RGC terminal division, the average trajectory ±S.D. and the average trajectory in WT situation.

We thus asked whether these apical MTs are important for RGC displacement using the MT depolymerizing drug colcemid. The drug was added at 32 hpf and Zn5 antibody staining was performed 16 hours later. The experiment revealed that many RGCs were retained at mid-retinal positions in drug-treated embryos (Fig. 4 F), while control RGCs formed a compact basal layer (Fig. 4 E). This suggested that MT depolymerization indeed interferes with RGC movement. To corroborate this finding and interfere with MT stability in single cells *in vivo*, we generated a construct to overexpress the MT destabilizing protein Stathmin 1 (Jourdain et al., 1997) under the control of heat shock promoter (*hsp70*) to obtain temporal control. We confirmed that Stathmin overexpression led to MT depolymerization in the *Tg(bactin:GFP-DCX)* line (Fig. S4 A). Embryos were injected with a *hsp70:Stathmin1-mKate2* plasmid, heat shocked at 24 hpf (to observe progenitor behavior) or 30 hpf (to observe RGC behavior) and imaged starting 4 hours after heat shock. Stathmin overexpression had no effect on progenitor cell behavior. These cells kept their bipolar morphology and interkinetic nuclear migration still occurred (Fig. S4 D). On the other hand, RGCs lost their basal process after apical division (while keeping the apical process) initially leading to a compromised movement (Fig. 4 G, n=22 N=5, Fig. 5 B). This initial loss of the basal process could be due to the fact that less MTs are present here and that these MTs were not acetylated (Fig. 4 C). In many cases, movement of RGCs was less directional than in the control situation as movement back towards the apical side often followed initial basal movement so that the net displacement of cells was marginal (Fig. S4 E and F, before apical process loss). This behavior could stem from the fact that apical MTs usually block nuclear movements back to apical positions. The strength of the observed phenotype correlated with the strength of Stathmin overexpression (data not shown).

**Figure 5.**
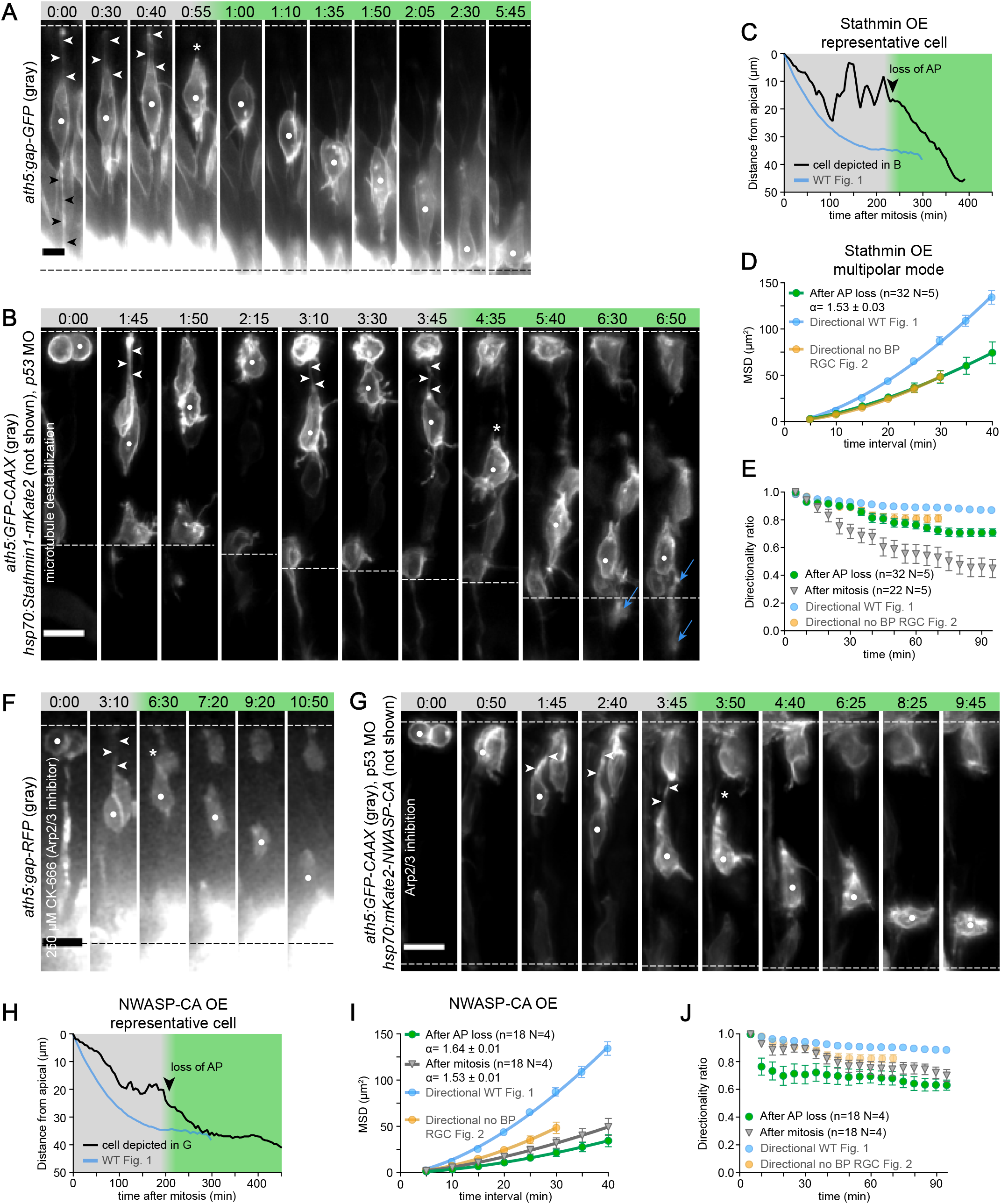
RGCs can switch to a multipolar migratory mode. A) A rare example of multipolar migration in control embryos. Gray phase: cell still has the basal and apical process. Green phase: directional multipolar mode. Dashed lines delimit the apical and basal sides of the retina. White dot: RGC followed. Arrowheads: apical and basal process. Asterisk: loss of apical process. Time is shown in hh:mm. Scale bar: 5 μm. B) Multipolar migration induced by microtubule destabilization. Stathmin1 overexpression was induced at 30 hpf. Movie starts at 34 hpf. Note higher protrusive activity upon loss of basal and apical process. Gray phase: cell still has the apical process. Green phase: directional multipolar mode. Dashed lines delimit the apical and basal sides of the retina. White dot: RGC followed. Arrowheads: apical process. Asterisk: loss of apical process. Blue arrow: axon. Time is shown in hh:mm. Scale bar: 10 μ m. C) Typical trajectory of RGC with destabilized microtubules from the montage in (B). Arrowhead: loss of the apical process (AP). For more trajectories see Fig. S4 E. The average WT trajectory is shown for comparison. D) MSDs of RGCs in multipolar migratory mode. The values are taken from the first 95 min after apical process loss. WT from Fig. 1 E and RGC without basal process from Fig. 2 G are plotted for comparison. E) Directionality ratios before and after apical process loss. The values are taken from the first 95 min after mitosis and the first 95 min after apical process loss. Directionality ratio at the end of the trajectory: After AP loss=0.71 After mitosis=0.45. WT from Fig. 1 F and RGC without BP from Fig. 2 J are plotted for comparison. F) Multipolar migration induced by Arp2/3 inhibition. The *ath5:gap-RFP* fish were imaged at a spinning disk microscope from 34 hpf. CK-666 was added at the start of imaging. Images were denoised in Fiji (ROF denoise). The dashed line delimits the apical and basal side of the retina. White dot: RGC followed. Arrowheads: apical process. Asterisk: loss of apical process. Time is shown in hh:mm. Scale bar: 10 μm. G) Multipolar migration induced by Arp2/3 inhibition. NWASP-CA overexpression was induced at 30 hpf. Movie starts at 34 hpf. Gray phase: cell still has the apical process. Green phase: directional multipolar mode. Dashed lines delimit the apical and basal sides of the retina. White dot: RGC followed. Arrowheads: apical process. Asterisk: loss of apical process. Time is shown in hh:mm. Scale bar: 10 μm. H) Typical trajectory of RGC after Arp2/3 inhibition (NWASP-CA overexpression) from the montage in (G). Arrowhead: loss of the apical process (AP). For more trajectories see Fig. S4 H. The average WT trajectory is shown for comparison. I) MSDs of RGCs after Arp2/3 inhibition (NWASP-CA overexpression). The values are taken from the first 95 min after mitosis and the first 95 min after apical process loss. WT from Fig. 1 E and RGC without basal process from Fig. 2 G are plotted for comparison. J) Directionality ratios of RGCs after mitosis and after apical process loss in the Arp2/3 inhibition condition (NWASP-CA overexpression). The values are taken from the first 95 min after mitosis and the first 95 min after apical process loss. Directionality ratio at the end of the trajectory: After AP loss=0.63 After mitosis=0.70. WT from Fig. 1 F and RGC without BP from Fig. 2 J are plotted for comparison.

In conclusion, these data show that progenitors and RGCs differ in their MT arrangements. RGCs contain stabilized MTs in the apical process when translocating basally and the stabilized MTs aid efficient RGC displacement.

### RGCs can switch to a multipolar migratory mode upon disturbance of MTs or Arp2/3 inhibition

We showed that stabilized MTs in the apical process and maintenance of the attachment of the basal process to the ECM are important factors for efficient RGC translocation. However, in both conditions we noted that, although a significant number of cells were observed at ectopic locations, many RGCs still reached the basal side (Fig. 3, C and D and Fig. 4 F). We thus asked how and when RGC movement took place in case the MT cytoskeleton or basal process attachment was compromised. In the control scenario, RGCs were observed to lose their basal process in very rare cases. These cells subsequently detached their apical process, increased their protrusive activity and moved basally in a multipolar migratory mode (Fig. 5 A). To test whether a comparable switch to multipolar migration could also be observed for RGCs upon interference with MT stability, we imaged Stathmin overexpressing cells for 12–16 hours. In this experimental setup Stathmin overexpression led to depletion of acetylated microtubules as shown by antibody staining (Fig. S4 B). As noted before, cells initially kept their apical process and did not generate much net movement towards basal positions (Fig. 4 G). Upon continued imaging, however, cells lost their apical attachment about 200 min after apical division (n=20 N=5, median). After that point, these cells switched to a multipolar migratory mode (Fig. 5, B and C) as seen for the rare control examples. These multipolar cells lost polarity, as shown by random positioning of the MT-organizing center (Fig. S4 C). Once multipolar migration started, instantaneous velocities increased (Fig. S4 E and F). Multipolar migration was directionally persistent, as shown by supralinear MSDs of the multipolar mode with an α value of 1.53±0.03 (Fig. 5 D). However, it was not as efficient as somal translocation of RGCs inheriting the basal process (Fig. 5 D) as also seen in the directionality ratio (Fig. 5 E). Similar observations were made upon Arp2/3 inhibition by CK-666 (Fig. 5 F). Upon drug treatment, RGCs first lost their basal process and later their apical process and translocated via the multipolar mode (Fig. 5 F). Also after overexpressing NWASP-CA we observed RGCs that fell into the multipolar mode of translocation to reach basal positions (Fig. 5 G, n=18 N=4), while this treatment had no effect on progenitors (Figure S4 G). However, in this scenario, the multipolar mode was not as directed as seen in the Stathmin overexpression condition (Fig. 5 H–J, Fig. S4 H and I). This could indicate that Arp2/3 activity is also involved in this mode of migration and not only for basal process attachment. Multipolar movement was also described previously for amacrine cells (ACs) (Chow et al., 2015). Therefore, we performed high time resolution imaging of translocating ACs to compare their behavior to that of RGCs. ACs also initially showed a bipolar morphology and moved only slowly towards the inner nuclear layer (Fig. S4 J). However, once these cells lost their apical process attachment, their velocities and directionality increased (Fig. S4, K–M) similarly to RGCs.

Collectively, these results indicate that RGC migration is a robust process and cells can adopt an alternative mode of movement when translocation depending on the presence of the basal process is impaired. Notably, this second mode of translocation is less efficient. Nevertheless, RGCs reach the basal retinal layer where they can polarize.

### Complete arrest of RGC translocation and their ectopic maturation disturbs retinal lamination

The surprising finding that RGC translocation can also occur via a multipolar migratory mode argued that RGC movement is particularly robust. We speculated that the robustness of RGC translocation exists, because a failure of RGC displacement would have severe consequences for retinal development. To test this, we needed a condition in which both displacement modes, the one depending on basal process attachment and MTs as well as the multipolar mode, were impaired. We noted that Rockout treatment led to very little disturbance of the apical actin pool (Fig. S3, E and F) and translocating RGCs kept their apical process over the whole time span of imaging up to 9.5 hrs (Fig. 3 A). However, we did not favor this condition, as analysis would involve spinning disk microscopy, which would distort experimental kinetics. We preferred a genetic condition instead. It was previously shown that overexpression of a membrane-targeted atypical protein kinase C zeta (aPKC-CAAX), led to restructuring of actin organization in progenitor cells (Strzyz et al., 2015). Therefore, we tested whether overexpression of this construct could suffice to block RGC basal process attachment but allow cells to keep their apical process. Phalloidin staining of the apical surface of corneal cells in the *Tg(hsp70:mKate2-aPKC-CAAX)* line revealed disorganized and less abundant microridges but increased cortical actin presence (Fig. S3 F). We then imaged this transgenic line in combination with a *Tg(ath5:gap-GFP)* line. Heat shock was performed between 30 hpf and 32 hpf so that aPKC-CAAX was present when RGC emergence was at its peak. Live imaging revealed that in this condition RGCs indeed lost their basal process after apical division and for more than 6 hours did not lose their apical process (while the apical process was lost after 200 min (median) in the stathmin condition). This resulted in RGCs projecting axons while being localized in apical and central regions of the retina. These cells never moved to the basal retinal layer (Fig. 6 A). The analogous phenotype was observed when aPKC-CAAX expression was restricted to the *ath5* lineage using *ath5:mKate2-aPKC-CAAX*. Also in this case cells after terminal division did not reach the basal retinal layer and extended an axon at central retinal positions (Fig. 6 D). Results from live imaging were corroborated by Zn5 staining, which showed that in the heat shock condition a significant portion of RGCs remained in mid-retinal regions (Fig. 6, B and C and Fig. S5 B).

**Figure 6.**
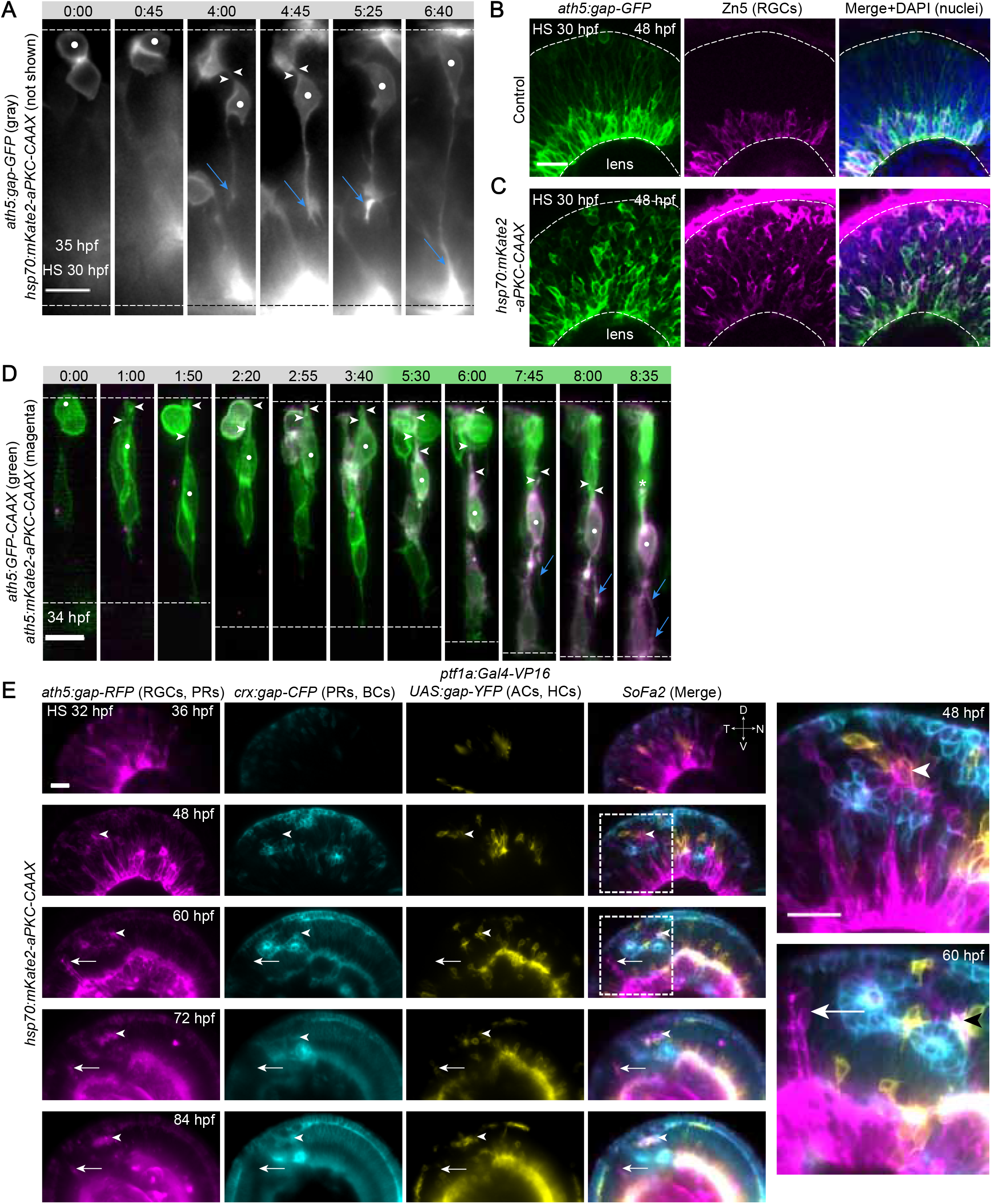
RGC translocation is stalled upon aPKC-CAAX overexpression and is not rescued over time. A) No RGC translocation upon aPKC-CAAX overexpression. Time is shown in hh:mm. Dashed lines delimit the apical and basal sides of the retina. White dot: RGC followed. Arrowheads: AP. Blue arrow: axon. Scale bar: 10 μm. B) Staining for differentiated RGCs with Zn5 antibody in control retina at 48 hpf. Note the basal RGC layer co-labeled with ath5:gap-GFP and Zn5. The dashed lines mark the apical and basal sides of the retina. Scale bar: 20 μm. C) aPKC-CAAX-overexpressing retina stained with Zn5 antibody at 48 hpf. Note disorganized RGC layer and ectopically differentiated RGCs. The strong signal beyond the apical side of the retina on the embryo surface is a nonspecific staining. D) aPKC-CAAX expression specifically in the *ath5* lineage stops translocation to the basal side and RGCs polarize at ectopic locations. Time is shown in hh:mm. Dashed lines delimit the apical and basal sides of the retina. White dot: RGC followed. Arrowheads: apical process. Asterisk: loss of apical process. Blue arrow: axon. Scale bar: 10 μm. E) Following ectopic RGCs in one embryo over time. The *SoFa2* transgenic fish (combination of *ath5:gap-RFP* (labeling RGCs and photoreceptors), *crx:gap-CFP* (labeling photoreceptors and bipolar cells), *ptf1a:Gal4-VP16 UAS:gap-YFP* (labeling horizontal cells and amacrine cells) was imaged every 12 hours and kept in the incubator between the time points. The ectopic RGCs developed on the left (temporal, T) side of the retina. The right (nasal, N) side developed as control (see Fig. S5 A) due to earlier differentiation there. Arrowhead: cluster of RGCs that trigger ectopic lamination of other cells types. Arrow: cluster of RGCs that interrupts the normal lamination without triggering ectopic layers. Dashed box: the magnified area. Scale bar: 20 μm.

To assess whether this impairment of RGC translocation had long-lasting effects on retinal lamination, we performed long-term imaging of *Tg(hsp70:mKate2-aPKC-CAAX)* fish crossed with the *SoFa2* line (Almeida et al., 2014), which labels membranes of all retinal cells using a combination of three fluorescent proteins. Fish were imaged in the LSFM every 12 hours between 36 hpf and 84 hpf to follow retinal lamination (Fig. 6 D). Control *SoFa2* retinas showed normal lamination (Fig. S5 A), whereas in combination with the *Tg(hsp70:mKate2-aPKC-CAAX)* line heat shocked at 32 hpf, RGCs were misplaced and this displacement was not corrected over development (arrows in Fig. 6 E). Notably, in this case the formation of other neuronal layers was also perturbed, with neurons aberrantly positioned and, in some cases, arranging around ectopically placed RGCs. This misplaced layering persisted until 96 hpf, and bipolar cells could sometimes be seen wrapped around ectopic RGC patches (Fig. 7, A and B). Lamination defects also occurred after mosaic injection of *ath5:mKate2-aPKC-CAAX* DNA (Fig. 7 C). However, the phenotypes were less frequent and milder in this condition because only a subset of RGCs expressed the construct and experienced translocation defects.

**Figure 7.**
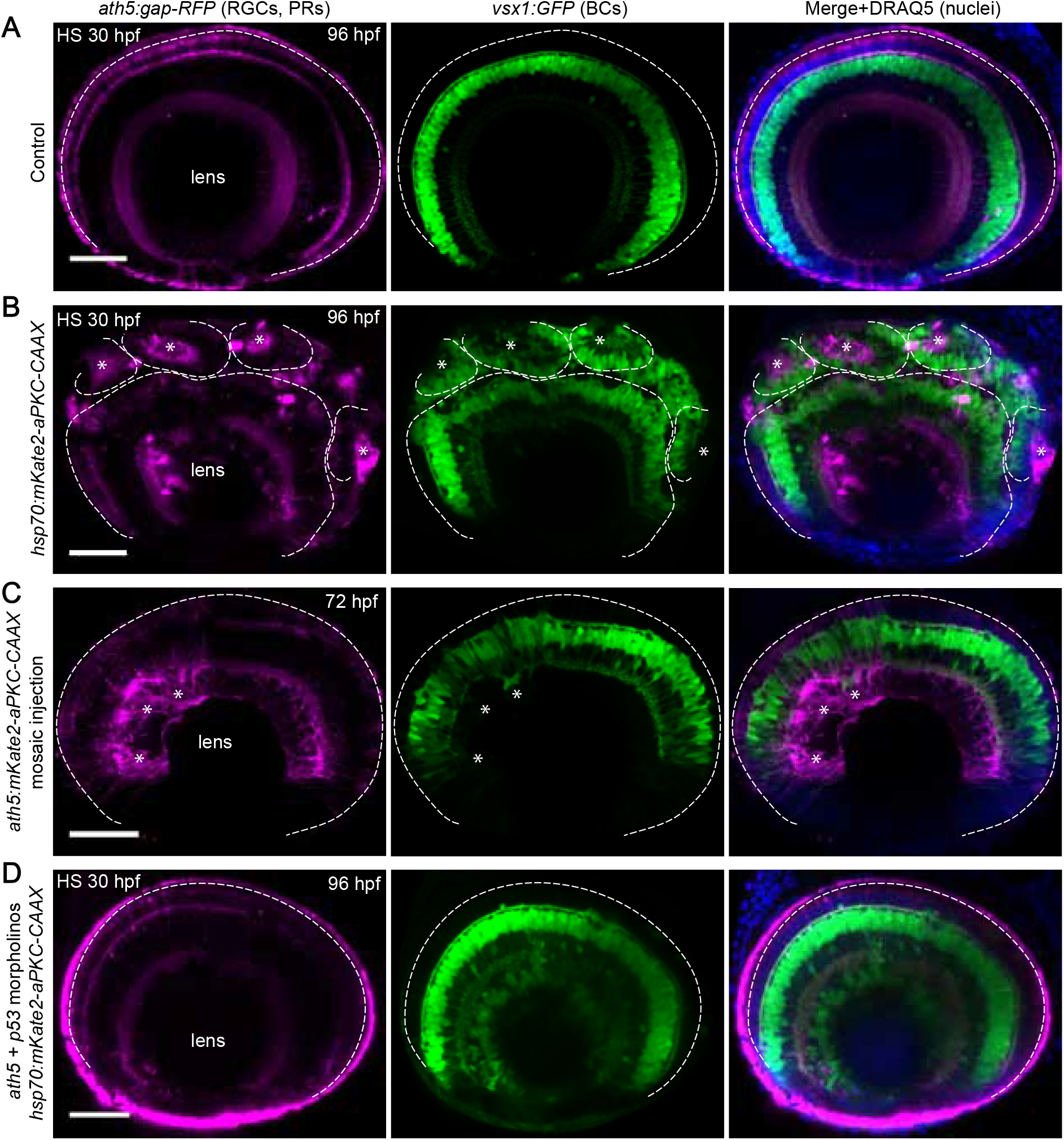
Ectopically differentiated RGCs induce retinal lamination defects. A) Control retina at 96 hpf with RGCs, photoreceptors (magenta) and bipolar cells (green) labeled. Additionally all nuclei were stained with DRAQ5 (blue). Dashed line delimits the apical side of the retina. Scale bar: 50 μm. B) Organizing role of ectopic RGCs. Asterisks: clusters of ectopic RGCs organizing the later born bipolar cells around themselves. Dashed lines delimit independent areas of lamination. Note the formation of inner plexiform layer-like structures in the areas of ectopic lamination. Scale bar: 50 μm. C) Lamination defects after mosaic expression of *ath5:mKate2-aPKC-CAAX*. Asterisks: area with the lamination defect, note the absence of defined inner plexiform layer. Scale bar: 50 μm. D) Retinal lamination upon aPKC-CAAX overexpression in the absence of RGCs. Note that the overall retina lamination is normal and the RGC layer is partially filled with bipolar cells. Showing that the disturbed lamination of bipolar cells in (B) is a non-cell autonomous effect caused by displaced RGCs. The *ath5:gap-RFP* reporter is still expressed in other cell types, but the RGCs are completely absent in *ath5* morphant as no optic nerve was formed. Scale bar: 50 μm.

We needed to ensure that the observed phenotypes in the *Tg(hsp70:mKate2-aPKC-CAAX)* line did not originate from defects in later translocation or neurogenesis events but were caused specifically by RGCs, which failed to translocate. To this end, we compared embryos heat shocked at different developmental stages. We overexpressed aPKC-CAAX at the peak of RGC generation, after the peak and when the RGC layer was fully formed, i.e. at 30 hpf, 36 hpf and 48 hpf respectively. Imaging at 96 hpf confirmed the severe lamination defect in the fish heat shocked at 30 hpf (Fig. S5 C). In contrast, fish heat shocked at 36 hpf showed only few ectopic cell clusters (arrowheads in Fig. S5 D) and fish heat shocked at 48 hpf (Fig. S5 E) were indistinguishable from controls (Fig. 7 A). Additionally, we used a published *ath5* morpholino (Pittman et al., 2008) to knock down this transcription factor as its absence is known to suppress the generation of RGCs (Kay et al., 2001). This enabled us to test whether lamination defects also occurred when no RGCs were present. *Tg(hsp70:mKate2-aPKC-CAAX, vsx1:GFP)* embryos injected with *ath5* morpholino, which were heat shocked at 30 hpf showed RGC layer partially filled with bipolar cells but did not develop an optic nerve (Fig. 7 D) as reported previously (Kay et al., 2001). The otherwise normal lamination (Fig. 7 D) of their retina at 96 hpf strongly argued that defects seen in the *Tg(hsp70:mKate2-aPKC-CAAX)* fish were indeed caused by the presence of ectopically differentiating RGCs. Thus, successful RGC translocation to basal positions is crucial for later retinal lamination events.

## Discussion

In this study, we investigate the first step of neuronal lamination in the zebrafish retina, the formation of the RGC layer. We reveal the kinetics of the RGC somal translocation mode and show that it represents a directed movement dependent on basal process attachment. When RGC somal translocation is impaired, cells can resort to a multipolar migratory mode. When both modes are inhibited though, RGCs fail to translocate and the resulting misplaced RGCs induce lamination defects, indicating that RGC translocation represents a crucial first step for the formation of other retinal layers. We summarize our main findings in (Fig. 8).

**Figure 8.**
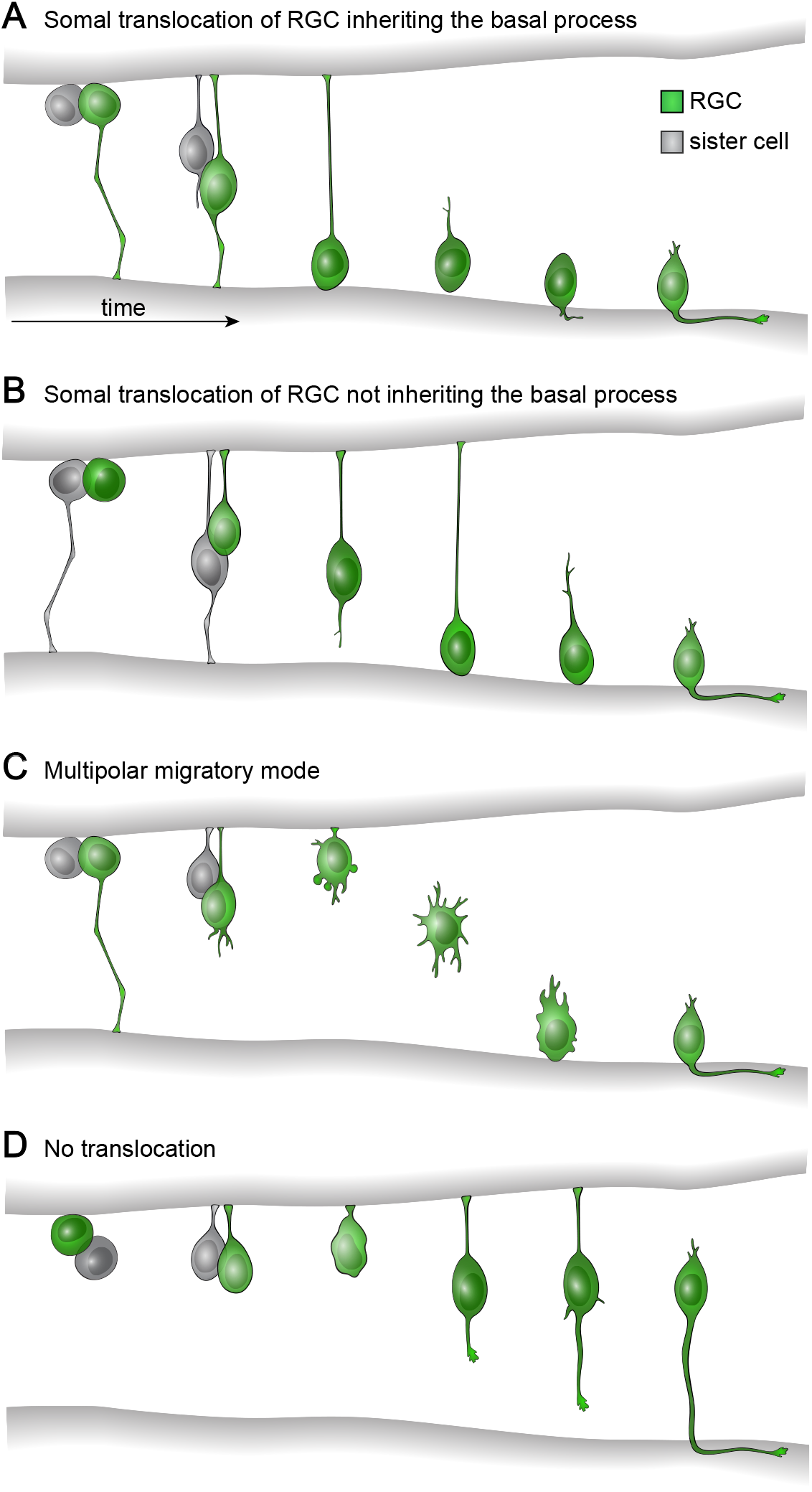
Summary of different RGC translocation scenarios. A) Somal translocation of RGC inheriting the basal process, a mode used by 80% cells. The RGC translocates basally faster than the sister cell. Directionally persistent somal translocation is followed by fine positioning, during which cells lose their apical process and eventually form the axon. B) Somal translocation of RGC not inheriting the basal process, a mode used by 20% of cells. The RGC initially lags behind the sister cell during the basal movement. Later it regrows the basal process and overtakes it. Translocation is less efficient than in A). Fine positioning phase, during which cells lose their apical process and eventually form the axon is shorter than in A). C) Multipolar migratory mode. This mode occurs after microtubule destabilization or Arp2/3 inhibition and in rare cases in control cells. After loss of basal process attachment, the RGC also detaches its apical process and increases protrusive activity. The cell then moves towards the basal side using the multipolar mode. This movement is less efficient than in A). Axon formation and RGC layer establishment are not affected by the multipolar movement. D) No translocation. In case RGC translocation is inhibited, cells are able to differentiate at ectopic locations instead of establishing a compact RGC layer, which has severe consequences for later retinal lamination.

### Similarities and differences between emerging RGCs and retinal progenitors

RGCs are the first neurons emerging in the vertebrate retina (Cajal, 1972; Nawrocki, 1985; Sidman, 1961) and at the time of their birth the majority of retinal cells are still apically dividing neuroepithelial progenitors. At first sight, apical progenitors and emerging RGCs appear similar: they both display bipolar morphology and the nucleus represents the bulkiest part of the cell. In addition, we found that both cell types translocate their nuclei towards basal positions more efficiently when they inherit the basal process. For both cell types, the basal translocation is fastest and most directional right after cell division, when the apical side of the retina represents a boundary preventing the movement back in apical direction (Leung et al., 2011). However, basal nuclear displacement is faster in RGCs, arguing that additional mechanisms are involved in these cells. Notably, the two cell types also differ in the organization of their MT cytoskeleton: in progenitor cells MTs are more dynamic, whereas RGCs feature stabilized MTs, which are involved in efficient basal translocation of their soma.

### Cellular requirements for RGC somal translocation

We show that the most efficient mode of RGC basal movement is directed somal translocation. Unlike many other neuronal migration events (Cooper, 2013), this movement is continuous instead of saltatory. Most RGCs inherit the basal process after their last apical division and this helps their efficient translocation. We show that when basal process attachment is impaired, RGC translocation is less efficient and cells often form their axons in mid-retinal positions. In addition to basal process attachment to the basal lamina, we show that MTs in the apical process are important for efficient basal translocation of RGCs. These stabilized MTs might prevent the nucleus from sliding back towards apical positions and allow the movement to be more persistent compared to progenitors, in which the nucleus often stalls or reverses apically. This notion is underlined by the finding that in the Stathmin overexpression condition, in which MTs are depolymerized, nuclei have less directional trajectories including jumps towards more apical positions.

It is possible that lateral interactions between RGCs and their neighboring progenitors, for example via contacts to their basal processes, play an inductive role in efficient somal translocation. Such observations have been made in other systems (Famulski et al., 2010; Elias et al., 2007; Marin et al., 2010) and it will be interesting to address whether and how neighboring cells influence RGC movements.

### RGCs can switch to a multipolar mode to reach basal positions when somal translocation is perturbed

Surprisingly, we noted that even when basal process attachment or stabilized MTs are disturbed, RGCs often still reached the basal RGC layer. In these scenarios, cells are stalled at the apical side of the retina before they retract their apical process and switch to a multipolar migratory mode. Once this multipolar migration is initiated, it is only slightly less efficient than the directional phase of somal translocation. The reason for the preservation of RGC directional persistence could lie in the fact that, when RGCs emerge, the majority of the retina is still occupied by stably attached bipolar progenitors. These tightly packed, elongated progenitors might prevent tangential movement of RGCs. It is also possible that additional guidance cues are involved in establishing directionality of multipolar migration and it will be interesting to explore these possibilities. Notably, RGCs are not the only neuronal cell type capable of employing two different translocation modes. It has been shown that projection neurons of the mouse cerebral cortex like RGCs can employ several different migratory modes including somal translocation and multipolar migration (Tabata and Nakajima, 2003; Noctor et al., 2004). We further showed that amacrine cells exhibit a similar pattern of slower bipolar phase of translocation with apical process and faster final multipolar movement after apical process loss. However, our result that amacrine cell translocation does not seem to be affected in the aPCK-CAAX ath5 morphant condition argues that the cascades driving apical detachment differ between these two cell types. This might not be surprising considering the fact that for amacrine cell translocation multipolar migration is the canonical mode of movement while for RGCs it is the exception. How apical detachment and the switch to multipolar migration differ between these two cell types will be an interesting point to solve in future studies.

This suggests that there are common patterns in neuronal migration in different organisms and that findings made in the zebrafish retina can be extrapolated to other areas of CNS development.

### RGC layer formation orchestrates subsequent retinal lamination events

The finding that RGCs can switch their mode of movement upon different perturbations reveals that RGC basal displacement is an unexpectedly robust process. We speculate that this robustness emerged to ensure later retinal lamination, because when RGC translocation is compromised, all following lamination events seem to be affected. Interestingly, we find that RGCs can organize later born cells around themselves when they are found at ectopic positions. It is described that RGCs play an organizing role during inner plexiform layer formation (Kay et al., 2004), but it is also known that the zebrafish retina can laminate relatively normally when RGCs are absent due to genetic depletion of the key transcription factors (Kay et al., 2001). This is also seen in the combined aPKC-CAAX overexpression *ath5* morphant condition in which retinal layering seems normal, most likely because no ectopic RGCs are present. Overall, this suggests that even though RGCs are not required for retinal lamination per se, when they are present, they play an organizing role regardless of their position.

We also observe that ectopic RGCs polarize normally and can form axons at any position in the retina, as reported previously (Zolessi et al., 2006; Randlett et al., 2011b). The time that has passed since the terminal mitosis, rather than progression of RGC translocation, seems to regulate when axonogenesis commences in a cell-autonomous manner and slower translocation of RGCs that did not inherit the basal process does not delay their axon outgrowth. Additionally, axonogenesis *in vivo* is distinct from neuronal behavior *in vitro* (Dotti et al., 1988), where a multipolar neuron grows several neurites, from which one is later selected to become the axon. In the retina, however, RGCs grow only one directed axon. This is not only true for cells that move via somal translocation, but also for cells that undergo multipolar migration, which is likely due to the polarizing cues present in the surrounding environment (Randlett et al., 2011b). Together, our findings indicate that RGCs are important to organize all later retinal development independently of their position. These cells may also have a potential to self assemble retinal structures in other tissue contexts, which might be an interesting entry road for therapeutic and transplantation approaches.

Beyond the findings on RGC translocation and retinal lamination events, this study exemplifies the importance of performing well controlled, *in toto* imaging to understand the kinetics and mechanisms of CNS development. Using the gentlest microscopy available allowed us to generate a comprehensive high quality dataset and study the process in question as close to the developmental ‘ground truth’ as currently possible (Stelzer, 2015). If we had limited our study to comparing the ‘before’ and ‘after’ situation, many important insights would have been missed, including the multipolar migration mode of RGCs and the importance of basal process inheritance. Obviously, such experiments are not feasible in every tissue or model system. Still, we would like to argue that studies like ours will generate important advances in the understanding of CNS formation and reveal general principles that can be extended to other, less accessible systems.

## Materials and Methods

### Zebrafish Husbandry

WT TL zebrafish were maintained and bred at 26°C. Embryos were raised in E3 medium at 28.5°C or 32°C and treated with 0.2 mM 1-phenyl-2-thiourea (PTU) (Sigma) from 8–10 hpf onwards to delay pigmentation. Medium was changed daily. All animal work was performed in accordance with European Union (EU) directive 2011/63/EU as well as the German Animal Welfare Act.

### Zebrafish transgenesis and transgenic lines

1 nl of Tol2 plasmid containing the ubiquitously expressed MT marker doublecortin *Tg(bactin:GFP-DCX)* or *Tg(hsp70:mKate2-aPKC-CAAX)* at 30 ng/μl and Tol2 transposase RNA at 50 ng/μl in ddH_2_O supplemented with 0.05% phenol red (Sigma) was injected into the cytoplasm of one-cell stage embryos. F_0_ embryos with fluorescence signal were grown to adulthood and Tg carriers were identified by outcross with WT fish. A list of Tg lines used can be found in the Supplemental Materials and Methods.

### Heat shock of embryos

To induce expression of the heat shock promoter (hsp70) driven constructs, the Petri dish with embryos was placed into a water bath set to 37°C or 39°C for 15–30 min depending on the construct. Imaging was started 3–4 hours after heat shock.

### Blastomere transplantations

Transplantations were performed as described previously (Randlett et al., 2013).

### DNA and morpholino injection

DNA constructs were injected into the cytoplasm of one-cell stage embryos. The DNA was diluted in ddH_2_O supplemented with 0.05% phenol red (Sigma) and the injected volume ranged from 0.5–1 nl. DNA concentrations ranged from 10–20 ng/μl and did not exceed 30 ng/μl even if multiple constructs were injected *(hsp70:EB3-mKate2*, 5 ng/μl). 0.5–1 ng of *laminin α1* morpholino (5′-TCATCCTCATCTCCATCATCGCTCA-3′) or 0.5–1 ng of control morpholino (5′-CCTCTTACCTCAGTTACAATTTATA-3′) were injected together with 2–4 ng *p53* morpholino (5′-GCGCAATTGCTTTGCAAGAATGT-3′), as reported previously (Randlett et al., 2011b). 2 ng of *ath5* morpholino (5′-TTCATGGCTCTTCAAAAAAGTCTCC-3′, all Gene Tools) was injected as reported previously (Pittman et al., 2008) to inhibit RGC formation. 2 ng of *p53* morpholino was also coinjected together with the *hsp70:Stathmin1-mKate2* and *hsp70:mKate2-NWASP-CA* constructs to alleviate the toxicity connected with construct overexpression.

### DNA constructs used and cloning strategies

Gateway cloning (Thermo Fisher Scientific) based on the Tol2 kit (Kwan et al., 2007) was used for all constructs. Details of cloning strategies and a list of all constructs used can be found in Supplemental Materials and Methods.

### Drug treatments

Colcemid (Enzo Life Sciences) was dissolved in DMSO as a 25 mM stock solution. The final concentration was 100 μM. CK-666 (Merck) was dissolved in DMSO as a 50 mM stock solution. The final concentration was 200 or 250 μM. Rockout (Santa Cruz) was dissolved in DMSO as a 50 mM stock solution. The final concentration was 50 or 100 μM. Aphidicolin (Sigma) was dissolved in DMSO as a 30 mM stock solution. The final concentration was 150 μM. Hydroxyurea (Sigma) was dissolved in H_2_O as a 1 M stock solution. The final concentration was 20 mM. All drug treatments were performed in 2 ml of E3 medium in a 24 well plate or a glass bottom dish. The same volumes of DMSO served as a control. The treatments started at the onset of RGC specification at around 32 hpf and lasted until 48 hpf, when embryos were fixed for immunostaining, or until the end of live imaging experiment.

### Immunofluorescence

All immunostainings were performed as whole mount on embryos fixed in 4% paraformaldehyde (Sigma) in PBS. The embryos were permeabilized with trypsin, blocked and incubated with the primary antibody for 3 days. The Zn5 antibody (ZIRC, RRID:AB_10013770) was used 1:50, the phosphorylated histone H3 antibody (Abcam, Cat# ab10543, RRID:AB_2295065) was used 1:500, the acetylated Tubulin antibody (Sigma-Aldrich Cat# T6793 RRID:AB_477585) was used 1:500, the Laminin α1 antibody (Sigma-Aldrich Cat# L9393, RRID:AB_477163) was used 1:250. Then the embryos were incubated for 3 days with an appropriate fluorescently labeled secondary antibody (Molecular Probes) 1:1000 and DAPI or 5 μM DRAQ5 (Thermo Fisher Scientific). Phalloidin-AlexaFluor 488 (Life technologies) was used 1:50. The acridine orange (Sigma) was dissolved at 2 μg/ml in E3 medium and live embryos were incubated in the solution for 30 min followed by 10 brief washes with E3 medium and imaging with the GFP filter set.

### Image Acquisition

#### Confocal Scans

Imaging was performed on LSM 510 or LSM 780 (Carl Zeiss Microscopy) using the Zeiss 40×/1.2 or 63×/1.2 C-Apochromat water immersion objective.

#### Time-lapse imaging Using LSFM

Imaging was performed as previously described (Icha et al., 2016) using the Lightsheet Z.1 (Carl Zeiss Microscopy). Briefly, the embryo was embedded in a 0.9% low melt agarose column and a 50–80 μm z stack of each eye was acquired in a single view, dual-sided illumination mode. Images were taken every 5 min for 12–16 hrs using the 10×/0.2 illumination objectives and a Plan-Apochromat 20×/1.0 W or 40×/1.0 W detection objective. The sample chamber was filled with E3 medium containing 0.01% MS-222 and 0.2 mM PTU and maintained at 28.5°C.

#### Time-lapse Imaging Using Spinning Disk Confocal Microscopy

Imaging was performed as described previously (Dzafic et al., 2015) with UPLSAPO 60×/1.3 silicon oil objective (Olympus).

#### Image Analysis

All the image data were processed in Fiji (Schindelin et al., 2012). The raw LSFM data were deconvolved in ZEN 2014 software (black edition, release version 9.0) using Nearest Neighbor algorithm. The drift was corrected with a manual drift correction plugin (http://imagej.net/Manual_drift_correction_plugin). The apoptotic cells were counted automatically with an ImageJ script (see Supplemental Materials and Methods). GraphPad Prism, version 6.0c for Mac OS (GraphPad Software) was used for statistical analysis and to create graphs.

#### Cell Trajectory, directionality ratio and MSD analysis

The translocation of cells was tracked manually in 2D in maximum projected substacks of the raw data by following the center of the cell body in Fiji using ImageJ plugin MtrackJ (Meijering et al., 2012). The resulting trajectories were analyzed as described previously (Norden et al., 2009; Leung et al., 2011) by calculating instantaneous velocities, MSDs and directionality ratios. MSDs and directionality ratios were calculated in the DiPer program (Gorelik and Gautreau, 2014) executed as a macro in Microsoft Excel. MSD values were fitted with 2D_Δ_t^α^ to estimate the α where D is the diffusion coefficient and _Δ_t is the interval between time points. The fit was done with non-linear regression (least squares) in GraphPad Prism.

## Author contributions

Conceptualization, C.N., J.I., Methodology, J.I., C.N., Investigation, J.I., C.G., M.R.-M., Writing – Original Draft, C.N., J.I. Writing – Review & Editing, C.N., J.I., Visualization, J.I., Supervision, C.N., J.I.

## Acknowledgements

We thank W. A. Harris, J. Mansfeld, I. K. Patten, and P. J. Strzyz for helpful comments on the manuscript. We also thank the Norden lab for useful discussions. We are grateful to C. Fröb, A. P. Ramos, S. Schneider, P. J. Strzyz and R. Swane, the light microscopy facility and the fish facility of the MPI-CBG for experimental help. We thank B. Lombardot from MPI-CBG Scientific computing facility image analysis plugins and to D. Sulcova for graphic works. We thank B. Ciruna, D. Gilmour, K. Kwan, L. LeClaire and J. Lippincott-Schwartz for sharing constructs. We would also like to thank the three anonymous reviewers for their constructive comments and suggestions that improved this manuscript.

J.I. is a member of the IMPRS of Cell, Developmental and Systems Biology. M.R.-M. was supported by CAPES/PDSE (99999.000424/2015-03). C.N. was supported by the Human Frontier Science Program (CDA-00007/2011) and the German Research Foundation (DFG) [SFB 655, A25].

The authors declare no conflict of interest.

## Supplemental materials and methods

### Transgenic lines used

| line | structures labeled | reference |
| --- | --- | --- |
| *Tg(bactin:GFP-DCX)* | all microtubules | this study |
| *Tg(hsp70:mKate2-aPKC-CAAX)* | aPKC in all membranes | this study |
| *Tg(bactin:ras-GFP)* | all membranes | gift from C.P. Heisenberg lab |
| *Tg(ath5:gap-GFP)* | membranes of RGCs and photoreceptors | (Zolessi et al., 2006) |
| *Tg(ath5:gap-RFP)* | membranes of RGCs and photoreceptors | (Zolessi et al., 2006) |
| *Tg(crx:gap-CFP)* | membranes of photoreceptors and bipolar cells | (Almeida et al., 2014) |
| *Tg(ptf1a:DsRed)* | amacrine and horizontal cells | (Jusuf et al., 2012) |
| *Tg(ptf1a:Gal4-VP16, UAS:gap-YFP)* | membranes of amacrine and horizontal cells | (Pisharath and Parsons, 2009)<br>(Williams et al., 2010) |
| *Tg(SoFa2)* | all retinal cells | (Almeida et al., 2014) |
| *Tg(vsx1:GFP)* | bipolar cells | (Kimura et al., 2008) |

### DNA constructs

*ath5:mKate2-aPKC-CAAX*

The coding sequence of mKate2-aPKC-CAAX (Strzyz et al., 2015) was assembled by classical cloning in the pCS2+ backbone and used as a template for creating the mKate2-aPKC-CAAX middle entry clone by PCR using Phusion polymerase (NEB) with primers:

F: 5′-ggggacaagtttgtacaaaaaagcaggctggATGGTGAGCGAGCTGATTAAGG-3′

R: 5′-ggggaccactttgtacaagaaagctgggtcTTAGGAGAGCACACACTTG-3′

The Ath5 promoter 5′ entry clone (Kwan et al., 2007) was combined with mKate2-aPKC-CAAX middle entry clone and pTol2+pA R4-R2 backbone (Villefranc et al., 2007).

*ath5:GFP-CAAX*

The Ath5 promoter 5′ entry clone (Kwan et al., 2007) was combined with GFP-CAAX middle entry clone (Kwan et al., 2007) and pTol2+pA R4-R2 backbone (Villefranc et al., 2007).

*ath5:GFP-DCX*

The coding sequence of human doublecortin GFP-DCX plasmid was a gift from Joseph Gleeson (Tanaka et al., 2004) (Addgene plasmid # 32852). The GFP-DCX middle entry clone was created by PCR using Phusion polymerase (NEB) with primers:

F: 5′-ggggacaagtttgtacaaaaaagcaggctggATGGTGAGCAAGGGCGAGG-3′

R: 5′-ggggaccactttgtacaagaaagctgggtcTTACATGGAATCACCAAGCG-3′

The GFP-DCX middle entry clone was combined with the Ath5 promoter 5′ entry clone (Kwan et al., 2007) and pTol2+pA R4-R2 backbone (Villefranc et al., 2007).

*ath5:H2B-RFP*

The H2B-RFP middle entry clone (Strzyz et al., 2015) was combined with Ath5 promoter 5′ entry clone (Kwan et al., 2007) and pTol2+pA R4-R2 backbone (Villefranc 2007).

*bactin:GFP-DCX*

The GFP-DCX middle entry clone (this study) was combined with the beta actin promoter 5′ entry clone (Kwan et al., 2007) and pTol2+pA R4-R2 backbone (Villefranc et al., 2007). The final construct was injected together with the Tol2 RNA to create the transgenic line.

*bactin: mKate2-ras*

The coding sequence of mKate2 with a membrane-targeting signal from Homo sapiens Harvey rat sarcoma viral oncogene homolog (HRAS) was used. The mKate2-ras middle entry clone was created by PCR using Phusion polymerase (NEB) with primers:

F: 5′-ggggacaagtttgtacaaaaaagcaggctggATGGTGAGCGAGCTGATTAAGG-3′

R: 5′-ggggaccactttgtacaagaaagctgggtcTCAGGAGAGCACACACTTGC-3′

The mKate2-ras middle entry clone (this study) was combined with the beta actin promoter 5′ entry clone (Kwan et al., 2007) and pTol2+pA R4-R2 backbone (Villefranc et al., 2007).

*hsp70:Arl13b-mKate2*

The coding sequence of human ADP-ribosylation factor-like protein 13B (Arl13b-mKate2) plasmid was a gift from B.Ciruna (University of Toronto). The Arl13b-mKate2 middle entry clone was created by PCR using Phusion polymerase (NEB) with primers:

F: 5′-ggggacaagtttgtacaaaaaagcaggctggATGTTCAGTCTGATGGCC-3′

R: 5′-ggggaccactttgtacaagaaagctgggtcTTATTTGTGCCCCAGTTT-3′

The Arl13b-mKate2 middle entry clone was combined with the hsp70 promoter 5′ entry clone (Kwan et al., 2007) and pTol2+pA R4-R2 backbone (Villefranc et al., 2007).

*pCS2+ Centrin-tdTomato*

This construct was a kind gift from D. Gilmour (EMBL).

*hsp70:EB3-mKate2*

To create the middle entry clone of human microtubule-associated protein RP/EB family member 3 (EB3, also called MAPRE3) tagged with mKate2 was amplified from pCS2+ EB3-mKate2 (Strzyz et al., 2015) by PCR using Phusion polymerase (NEB) with primers:

F: 5′-ggggacaagtttgtacaaaaaagcaggctggATGGCCGTCAATGTGTACTCC-3′

R: 5′-ggggaccactttgtacaagaaagctgggtcTCATCTGTGCCCCAGTTTGC-3′

The EB3-mKate2 middle entry clone was combined with the hsp70 promoter 5′ entry clone (Kwan et al., 2007) and pTol2+pA R4-R2 backbone (Villefranc et al., 2007).

*hsp70:GalT-RFP*

The GalT-RFP (N-terminal 61 amino acid fragment of human galactosyl transferase) plasmid was a gift from J. Lippincott-Schwartz (NIH). The GalT-RFP middle entry clone was created by PCR using Phusion polymerase (NEB) with primers:

F: 5′-ggggacaagtttgtacaaaaaagcaggctggATGAGGCTTCGGGAGCCGC-3′

R: 5′-ggggaccactttgtacaagaaagctgggtctTTAGGCGCCGGTGGAGTGG-3

The GalT-RFP middle entry clone was combined with the hsp70 promoter 5′ entry clone (Kwan et al., 2007) and pTol2+pA R4-R2 backbone (Villefranc et al., 2007).

*hsp70:mKate2-aPKC-CAAX*

The construct was published previously (Strzyz et al., 2015). It was injected together with the Tol2 RNA to create the transgenic line.

*hsp70:mKate2-NWASP-CA*

The coding sequence of zebrafish wasb gene was amplified from cDNA. The truncation was created based on homology with a previously published human construct (Rohatgi et al., 1999). The last 55 C-terminal amino acids act as dominant negative and inhibit Arp2/3 complex. The NWASP-CA 3′ entry clone was created by PCR using Phusion polymerase (NEB) with primers:

F: 5′-ggggacagctttcttgtacaaagtggctGTGTCTGAATCCCCGGAC-3′

R: 5′-ggggacaactttgtataataaagttgcTTAGTCATCCCATTCATCATCT-3′

The NWASP-CA 3′ entry clone was combined with the hsp70 promoter 5′ entry clone (Kwan et al., 2007), mKate2 middle entry clone (gift from A. Oates laboratory) and pTol2+pA R4-R3 backbone (Kwan et al., 2007).

*hsp70:PCNA-GFP*

The coding sequence of human proliferating cell nuclear antigene (PCNA) was used as a template. The EGFP-PCNA middle entry clone was created by PCR using Phusion polymerase (NEB) with primers:

F: 5′-ggggacaagtttgtacaaaaaagcaggctggATGGTGAGCAAGGGCGAGG-3′

R: 5′-ggggaccactttgtacaagaaagctgggtcCTAAGATCCTTCTTCATCCTCG-3′

The EGFP-PCNA middle entry clone was combined with the hsp70 promoter 5′ entry clone (Kwan et al., 2007) and pTol2+pA R4-R2 backbone (Villefranc et al., 2007).

*hsp70:Stathmin1-mKate2*

The construct was published previously (Taverna et al., 2016).

## Script for detection of apoptotic cells in the zebrafish

Author: Benoit Lombardot (Scientific Computing Facility, MPI-CBG)

Date: 2016-07-19

### Description

This script is able to count and characterize fluorescent object in a user-defined region of interest. In particular it was developed to count apoptotic cell cluster (as well as their size) in head region of zebrafish.

### Usage

This section describes how to use the script (i.e. how to perform the analysis and what kind of image it can analyze).

To use the script, download start Fiji, then go to the menu ***Help>Update…***. In the ImageJ Updater window that just open click ***manage update site***. A new window opens where one can check the ***3D ImageJ Suite*** update site. Finally close the windows and restart Fiji.

Once Fiji is started, open the script editor (menu File>New>Script) and copy the content of the code section in the editor window. In the menu ***language*** select ***python*** this should color the syntax of the script. Optionally you can save the disk to your hard-drive (menu ***File>Save***). The script is now ready to be run.

Open an image and draw a region of interest around the region to analyze. The image can be a 2d image or a z stack with multiple channels. For 3d images the regions of interest needs only to be defined in 2D on one slice of the stack. One of the channels should show objects to detect in fluorescence modality. The objects should be sparse in 3d images as the image will be projected to simplify the analysis and a high density of objects would cause some overlap and degrade the quality of the analysis. Once this is done you can run the script

Once the script starts, a window will open requesting some parameters (see Figure 1):

- **channel to analyse**: the number of the fluorescence channel
- **maximum object radius (in pixel)**: details larger will be filtered out of the image
- **minimum object area (in pixel)**: object smaller than that size will be discarded
- **use automatic threshold**: if checked an Otsu threshold will be used to determine the foreground object
- **manual threshold**: This the threshold value used in case the previous parameter is not checked
- **use watershed**: if checked stuck objects will be separated with the watershed algorithm

**Figure 1:**
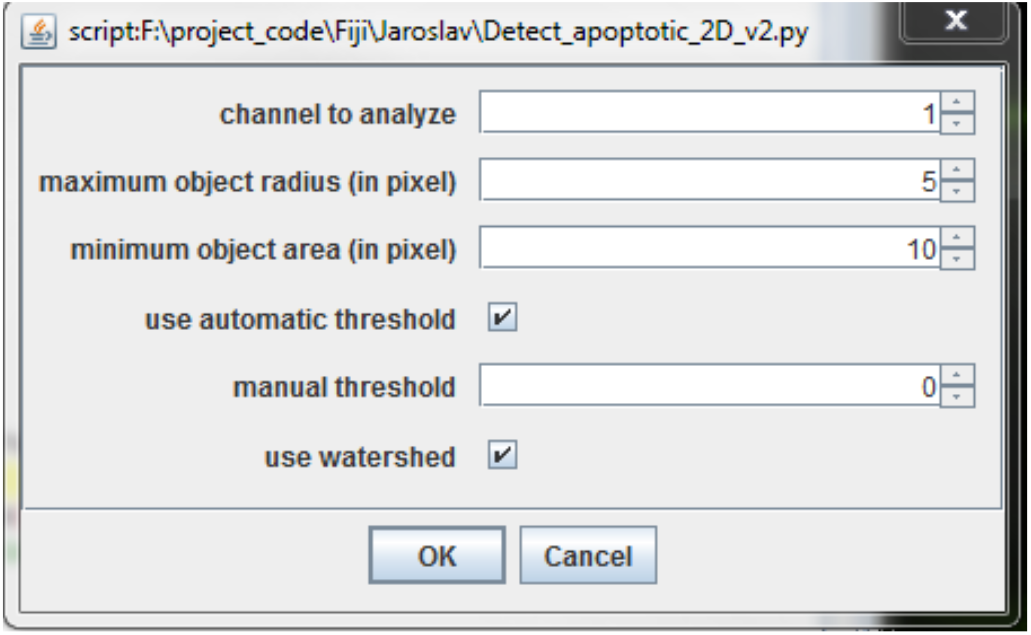
Window requesting parameters when the script is started. Once parameters are set, one can click ok and proceed with the analysis of the image. The plugin outputs a result image and some measures. The image shows the user-selected region of interest in green and detected objects contour in red (see Figure 2). The measure window indicates the number of objects their sum area (in image unit), their sum intensity and the area of the region analyzed (in image unit).

**Figure 2:**
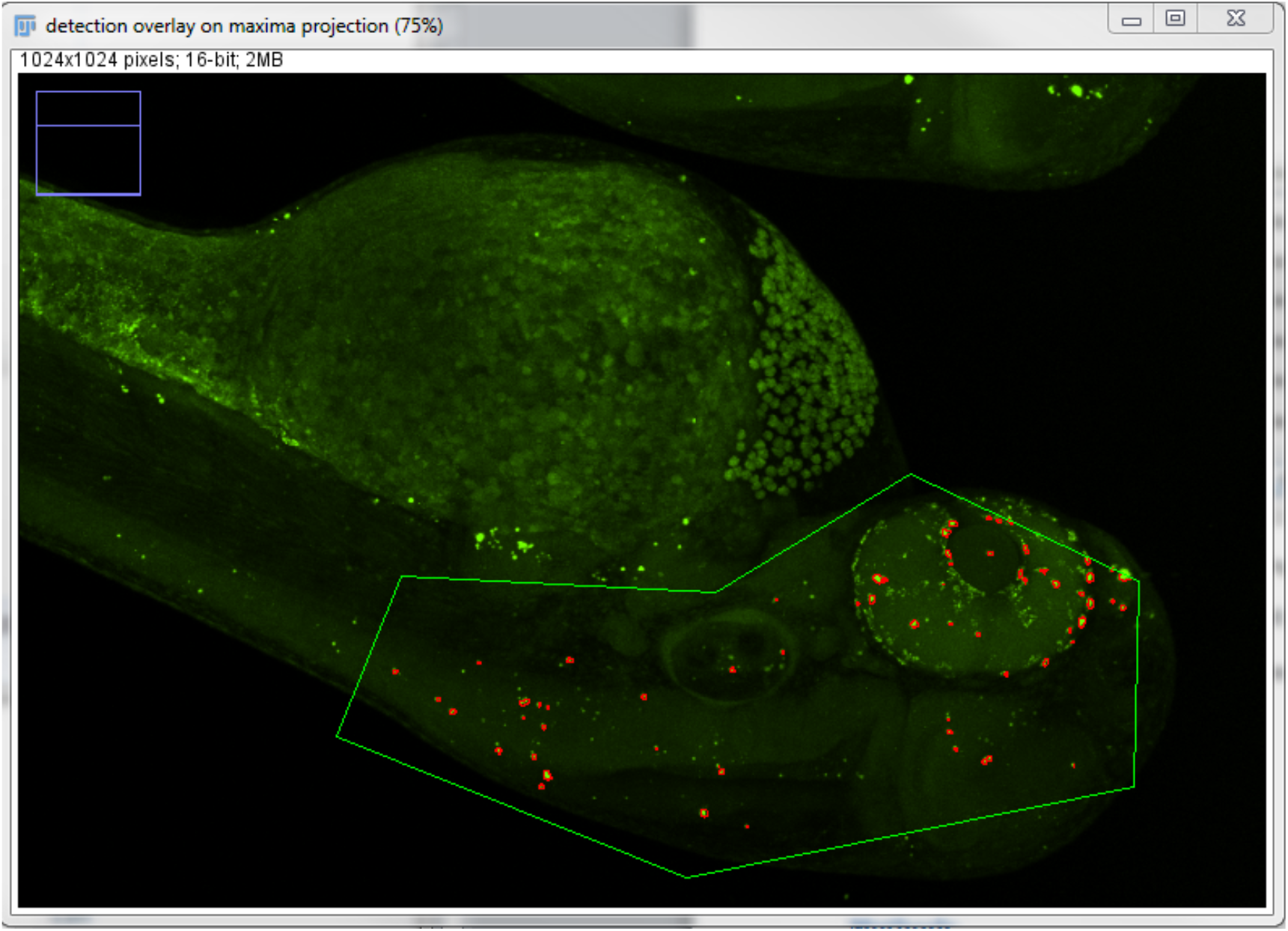
Result image displayed at the end of the script. The user-selected region of interest is shown in green and the detected objects are shown in red.

### Methods

This section details the image analysis strategy that was used to count the objects in the fluorescent channel.

The problem is first reduced to 2 dimensions to simplify the analysis. This is done by doing a maximum projection of the fluorescent channel along the z direction of the stack. The rolling ball algorithm is then applied to the resulting image to remove the background signal. The latter could be due for instance to tissue auto-fluorescence. The rolling ball radius is chosen larger than the radius of the object to be detected which can be estimated manually (for instance with Fiji) in the image. The result is then filtered with a Gaussian kernel of standard deviation 1 pixel to remove noise and regularize the image. The image is then thresholded with the Otsu method or manually if user requested it. In the case the user chose a manual threshold, it can be estimated from a few images by looking at the typical intensity of objects to detect and choosing a threshold worth 5 percent of that value. Then same manual threshold can then be used to analyze all the images of an experiment. After the thresholding one obtains a mask image showing the detected objects. If object are stuck together using the watershed option can help to separate the object to improve count quality. The objects outside the user-defined region of interest are then set to zero. The number of object in the final mask is then determined as the number of connected components in the image. Additionally the sum area and intensity of the objects are calculated as well as the region of interest that was analyzed.

### Code

~~~
#@int (label=“channel to analyze”)ch
#@int (label=“maximum object radius (in pixel)”)R
#@int (label=“minimum object area (in pixel)”)minArea
#@boolean (label=“use automatic threshold”)isAutoThresh

#@float (label=“manual threshold”)T
#@boolean (label=“use watershed”)isWatershed
# Author: Benoit Lombardot, Scientific Computing Facility, MPI-CBG
# 2014-06-26 first version for Jaroslav Icha (Norden Lab)
# 2016-07-19 simplify parameter input and update comments for publication (for Jaroslav Icha (Norden Lab))

# The code was tested with Fiji 2.0.0 rc 49 / 1.51d on July 19^th^ 2016

from ij import IJ;
from ij.gui import Roi;
from ij.plugin import Duplicator;
from ij.plugin import ZProjector as Zproj
from ij.process import ImageStatistics;
from ij.gui import ShapeRoi;
from ij import ImagePlus;
from ij.process import ByteProcessor;
from ij.measure import ResultsTable;
from ij import WindowManager;
from ij.gui import Overlay;
from ij import WindowManager;
from ij.text import TextWindow;

from java.awt import Color;

from util import FindConnectedRegions;

from mcib3d.image3d import ImageHandler;
from mcib3d.image3d.processing import FastFilters3D;
from mcib3d.image3d.regionGrowing import Watershed3D;

# grab visible image
imp0 = IJ.getImage();
stats_0 = imp0.getStatistics(ImageStatistics.AREA);
area_analyzed = stats_0.area;

# grab roi
roi = imp0.getRoi();
imp0.killRoi();

ratioXoverZ= imp0.getCalibration().pixelDepth/imp0.getCalibration().pixelWidth;
# duplicate the fluorescence channel
nZ = imp0.getNSlices();
nF = imp0.getNFrames();
imp = Duplicator().run(imp0, ch, ch, 1, nZ, 1, nF);

# max intensity projection zproj = Zproj(imp);
zproj.setMethod(Zproj.MAX_METHOD);
zproj.setStartSlice(1);
zproj.setStopSlice(nZ);
zproj.doProjection();
imp_proj = zproj.getProjection();

imp_proj_4disp = Duplicator().run(imp_proj);
# filter: gauss(1), bg removal, threshold (or maxima then watershed)
IJ.run(imp_proj, “Gaussian Blur…”, “sigma=1”);
IJ.run(imp_proj, “Subtract Background…”, “rolling=“+str(R) );

# set a threshold
if isAutoThresh:
         imp_proj.getProcessor().setAutoThreshold(“Otsu dark”);
         T = imp_proj.getProcessor().getMinThreshold();
else:
         stats = imp_proj.getStatistics(ImageStatistics.MIN_MAX);
         IJ.setThreshold(imp_proj,T,stats.max);
zproj2 = Zproj(imp);
zproj2.setMethod(Zproj.SUM_METHOD);
zproj2.setStartSlice(1);
zproj2.setStopSlice(nZ);
zproj2.doProjection();
imp_proj_4meas = zproj2.getProjection();

# performed image segmentation : output is a B&W image imp_proj
if isWatershed == True:
         img = ImageHandler.wrap(imp_proj);
         max_img = FastFilters3D.filterImage(img, FastFilters3D.MAXLOCAL, 3, 3, 3, 2, True);
         ws_img = Watershed3D(img, max_img, int(T), int(T) );
         label_imp = ImagePlus(“labelled peaks”,
ws_img.getWatershedImage3D().getImagePlus().getImageStack());
         # object are separated by pixel of value one. in 2d this work nicely with particle analyzer
         imp_proj = label_imp;
         stats = imp_proj.getStatistics(ImageStatistics.MIN_MAX);
         IJ.setTh reshold(imp_proj,2,stats.max);
else:
         IJ.run(imp_proj, “Convert to Mask”, “ “);

# Take the intersection of the manually inputed roi and detectected fluorescence spot
IJ.run(imp_proj, “Create Selection”, ““);
roi2 = imp_proj.getRoi();
imp_proj.killRoi()
if roi!=None:
         roi2.and(ShapeRoi(roi) );
imp_proj.setRoi(roi2);
IJ.run(imp_proj, “Create Mask”, “ “);
mask = IJ.getImage();

# analysis of the detected objects
imp_proj_4meas.setTitle(“imageToAnalyze”);
imp_proj_4meas.killRoi();
imp_proj_4meas.show();
IJ.run(“Set Measurements…”, “area mean redirect=“+ imp_proj_4meas.getTitle() +” decimal=3”);
IJ.run(mask, “Analyze Particles…”, “size=“+str(minArea)+”-Infinity display clear”);

imp_proj_4meas.hide();
imp_proj_4meas.flush();
mask.hide();
mask.flush();

# browse the value and count the object obove the threshold
rt = ResultsTable.getResultsTable();
frame = WindowManager.getFrame(“Results”);
if not frame==None:
         frame.close(False);
nval = rt.getCounter();
count = 0;
sumI = 0;
sumA = 0;
for i in range(nval):
         count = count+1;
         sumI = sumI + rt.getValue(“Mean”,i);
         sumA = sumA + rt.getValue(“Area”,i);

# display results
rt_summary = ResultsTable();
rt_summary.incrementCounter();
rt_summary.addValue(“count”,count);
rt_summary.addValue(“sum area”,sumA);
rt_summary.addValue(“sum intensity”,sumI);
rt_summary.addValue(“area analyzed”,area_analyzed);

#display result table
rt_summary.show(“image summary”);
# display overlay on the max projection
ov = Overlay();
roi2.setStrokeColor(Color. red);
ov.add(roi2);
if roi!=None:
         roi.setStrokeColor(Color.green);
         ov.add(roi);
         imp0.setRoi(roi);

imp_proj_4disp.setOverlay(ov);
imp_proj_4disp.setTitle(“detection overlay on maxima projection”)
imp_proj_4disp.show();
IJ.run(imp_proj_4disp, “Enhance Contrast”, “saturated=0.5”);
~~~

**Figure S1. Related to Figure 1.**
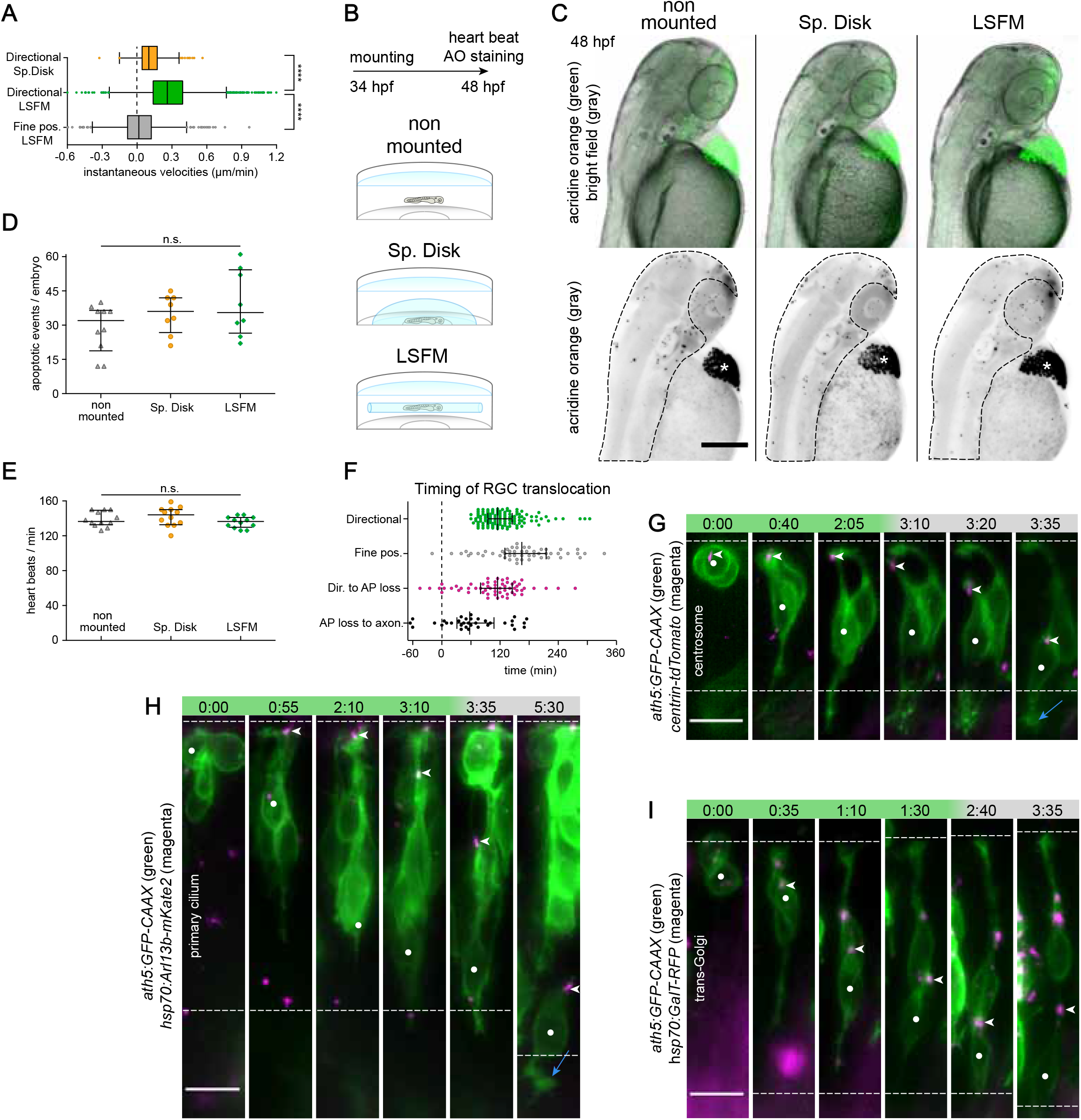
A) Comparison of instantaneous velocities during RGC directional phase between spinning disk confocal microscope and LSFM and between directional phase and fine positioning phase in LSFM. The values are taken from the first 95 min (or less) after mitosis or from the first 95 min of the fine positioning. The instantaneous velocities are calculated from the one-dimensional movement along the apico-basal axis of the retina. The movement from apical to basal has a positive sign; the reverse movement has a negative sign. Directional Sp. Disk (n=21 N=4, 399 data points), Directional LSFM (n=140 N=24, 2587 data points), Fine pos. LSFM (n=83 N=22, 1539 data points). 38 outliers were discarded by ROUT (Q=1.0%) for the plotting purposes, not for the statistical testing, which was done on the whole dataset. The differences between instantaneous velocities are statistically significant: Directional LSFM vs. Fine pos. LSFM Mann-Whitney U test p<0.0001; Directional LSFM vs. Directional Sp. Disk Mann-Whitney U test p<0.0001. The data are shown as Tukey boxplot (box shows median and interquartile range and whiskers show 1.5 of the interquartile range). The median instantaneous velocity Directional Sp. Disk= 0.101 μm/min, Directional LSFM =0.261 μm/min, Fine pos. LSFM=0.015 μm/min. B) Scheme of the experiment testing the influence of mounting for different microscopes on fish viability. The identical composition of E3 medium and agarose was used for both Sp. Disk and LSFM mounting and the dishes were kept together in an incubator overnight at 28.5°C. C) Acridine orange staining for apoptotic cells. The maximum intensity projection is shown for acridine orange staining. The bright field image was created with extended depth of field plugin in Fiji. Dashed area: area of apoptotic events count. Asterisk: hatching gland. Scale bar: 200 μm. D) Number of apoptotic events does not differ among the mounting strategies. N=10, 8 and 8 fish respectively. Bars represent median and interquartile range. Median non-mounted=32 events, median Sp. Disk=36 events, median LSFM=36 events. ANOVA p=0.1458. E) Heart rate does not differ among the mounting strategies. Heart beats were counted manually for 20 s in 12 fish for each condition. Bars represent median and interquartile range. Median non-mounted=137 beats/min, median Sp. Disk=144 beats/min, median LSFM=137 beats/min. ANOVA p=0.2149. F) Timing of key events during RGC translocation. The duration of directional phase: median 115 min (n=140 N=24). The duration of fine positioning phase, i.e., the time from the end of directional movement to axonogenesis: median 165 min (n=50 N=19). The time from the end of directional phase to apical process (AP) loss: median 115 min (n=58 N=21). The time from apical process loss to axonogenesis: median 57.5 min (n=33 N=16). The whiskers show median and interquartile range. G) Centrosome position during RGC translocation. *centrin-tdTomato* DNA was injected to label centrosome together with *ath5:GFP-CAAX*. H) Primary cilium position during RGC translocation. *hsp70:Arl13b-mKate2* DNA was injected to label primary cilium together with *ath5:GFP-CAAX*. I) Golgi apparatus position during RGC translocation. Galactosyl transferase fragment *hsp70:GalT-RFP* DNA was injected to label trans-Golgi together with *ath5:GFP-CAAX*. G), H), I) Green phase: directional movement. Gray phase: fine positioning. Dashed lines delimit the apical and basal sides of the retina. White dot: RGC followed. Arrowheads: organelle followed. Blue arrow: axon. Time is shown in hh:mm. Scale bar: 10 μm.

**Figure S2. Related to Figure 2.**
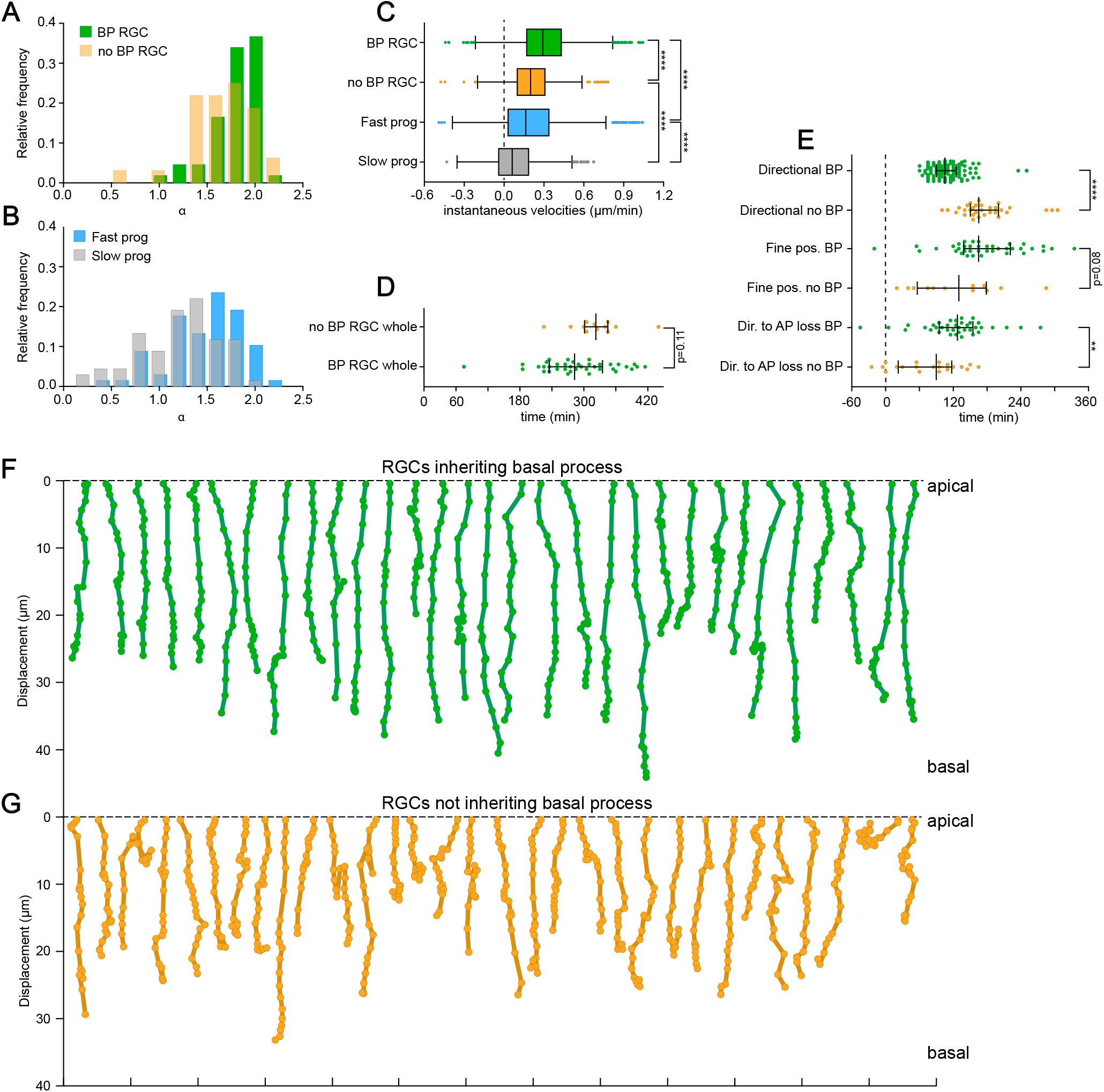
A) Alpha value distribution from MSDs of RGCs inheriting and not inheriting the basal process in directional phase (data from Fig. 2 G). Median BP RGCs=1.83, no BP RGCs=1.70. The differences between alpha values are statistically significant: BP RGCs vs. no BP RGCs Mann-Whitney U test p=0.03. B) Alpha value distribution from MSDs of fast and slow progenitor nuclei after mitosis (data from Fig. 2 H). Median Fast prog=1.53, Slow prog=1.26. The differences between alpha values are statistically significant: Fast progs vs. Slow progs Wilcoxon matched-pairs signed rank test p<0.0001. C) Comparison of instantaneous velocities among cells inheriting and not inheriting the basal process in directional phase (after mitosis). The values are taken from the first 70 min after mitosis. The instantaneous velocities are calculated from the one-dimensional movement along the apico-basal axis of the retina. The movement from apical to basal has a positive sign; the reverse movement has a negative sign. BP RGC (n=109 N=23, 1526 data points), no BP RGC (n=31 N=19, 448 data points), Fast progenitor (n=68 N=7, 952 data points), Slow progenitor (n=68 N=7, 952 data points). 45 outliers were discarded by ROUT (Q=1.0%) for the plotting purposes, not for the statistical testing, which was done on the whole dataset. The differences between instantaneous velocities are statistically significant: BP RGCs vs. no BP RGCs Mann-Whitney U test p<0.0001; Fast progenitors vs. Slow progenitors Wilcoxon matched-pairs signed rank test p<0.0001; no BP RGCs vs. Slow progenitors Mann-Whitney U test p<0.0001; BP RGCs vs. Fast progenitors Mann-Whitney U test p<0.0001. The data are shown as Tukey boxplot (box shows median and interquartile range and whiskers show 1.5 of the interquartile range). The median BP RGCs=0.29 μm/min, no BP RGCs=0.20 μm/min, Fast progenitors=0.16 μm/min, Slow progenitors=0.06 μm/min. D) Timing of the whole RGC translocation event (from mitosis to axonogenesis) in RGCs with and without basal process. BP median=282.5 min (n=38 N=18), No BP median=322.5 min (n=12 N=9), Mann-Whitney U test p=0.11. The median and interquartile range is shown. E) Timing of events during RGC translocation depending on the basal process inheritance. Median values: Directional BP=105 min (n=109 N=23), Directional no BP=165 min (n=31 N=19), Mann-Whitney U test p<0.0001. Median values: Fine positioning BP=165 min (n=38 N=18), Fine positioning no BP=130 min (n=12 N=9) Mann-Whitney U test p=0.08. Median values: From end of directional phase to apical process loss BP=127.5 min (n=38 N=16), no BP=90 min (n=20 N=15), Mann-Whitney U test p=0.003. The median and interquartile range is shown. F) Trajectories of RGCs inheriting the basal process. The representative 2D trajectories of 31 RGCs inheriting the basal process were plotted as manually tracked over the first 95 min after the terminal division. G) Trajectories of RGCs not inheriting the basal process. The 2D trajectories of all 31 RGCs not inheriting the BP were plotted as manually tracked over the first 95 min after the terminal division.

**Figure S3. Related to Figure 3, 5 and 6.**
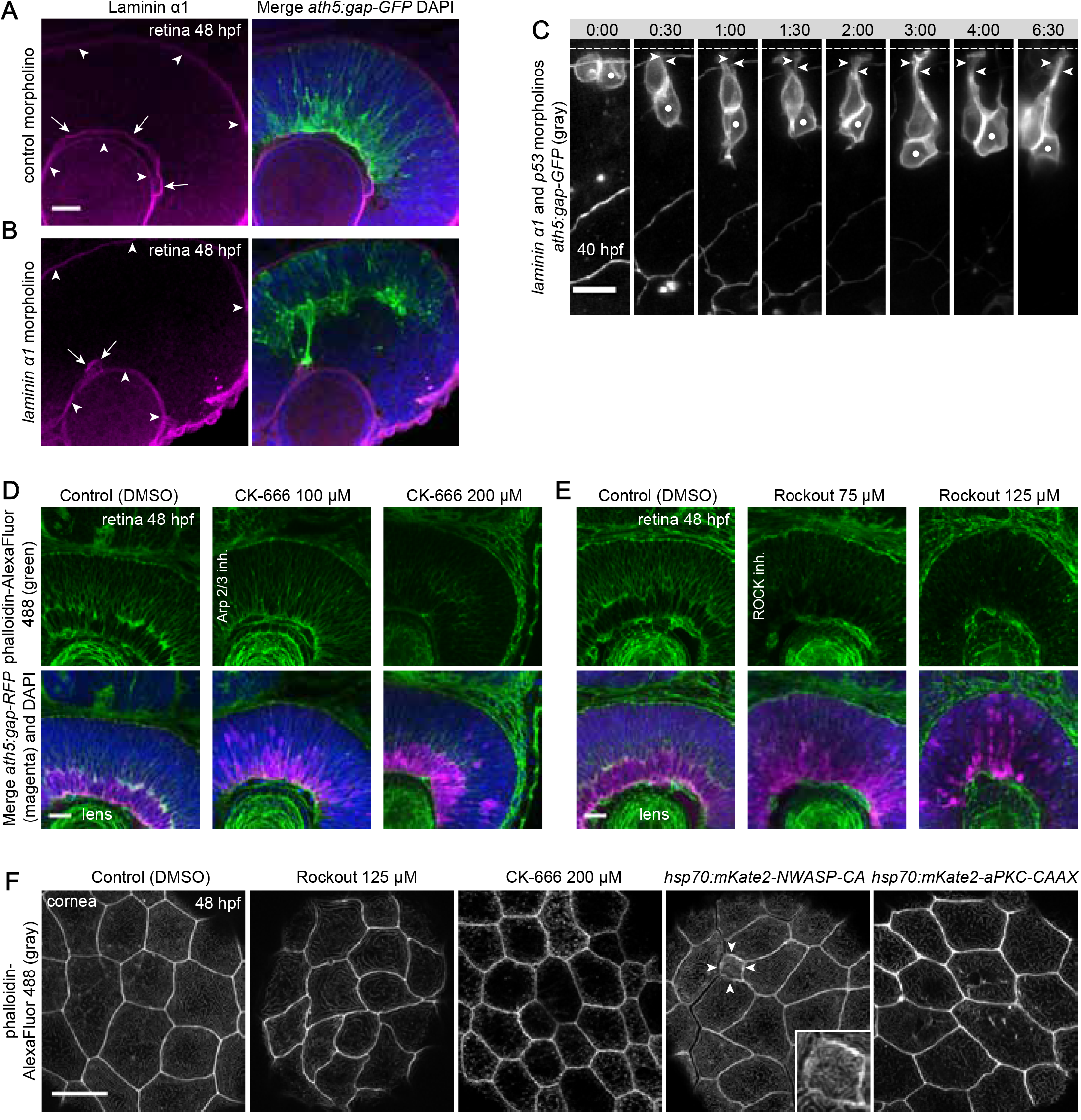
A) Laminin α1 distribution in control retina at 48 hpf. Laminin channel was denoised in Fiji (ROF denoise). Note the three distinct layers of Laminin (magenta) in the developing eye and the RGCs (green) forming a basal layer of the retina. Arrows: Laminin in retinal basal membrane Arrowheads: Laminin in lens and basal membrane of the retinal pigment epithelium. Scale bar: 20 μm. B) Laminin α1 distribution after Laminin knockdown. Note that Laminin is still present in the lens and basal membrane of the retinal pigment epithelium but mostly absent in the retinal basal membrane causing a tissue-wide lamination problem with RGC layer forming in the central retina. Arrows: Laminin in retina basal membrane. Arrowheads: Laminin in lens and basal membrane of the retinal pigment epithelium. Note that axons of RGCs associate with a small patch of Laminin left in the retinal basal membrane. C) RGC translocation is perturbed after Laminin α1 knockdown. Note that after division the cells lose the basal process and do not translocate basally. Dashed lines delimit the apical side of the retina. White dot: RGC followed. Arrowheads: apical process. Time is shown in hh:mm. Scale bar: 10 μm. D) Effect of Arp 2/3 inhibition on actin cytoskeleton in the retina. Note the decreased actin staining in the 200 μM condition in the basal and central retina. The apical actin belt associated with adherens junctions is less affected. The RGC layer labeled by *ath5:gap-RFP* is disorganized. Scale bar: 20 μm. E) Effect of ROCK inhibition on actin cytoskeleton in the retina. Note the decreased actin staining in the 75 μM and 125 μM condition in the basal and central retina. The apical actin belt associated with adherens junctions is less affected. The RGC layer labeled by *ath5:gap-RFP* is disorganized. Scale bar: 20 μm. F) Staining of apical actin by phalloidin in corneal cells at 48 hpf. Actin is present as two structures: it is associated with adherens junctions around the cell membranes and in microridges all over the apical membrane. The embryos were incubated with the inhibitors from 34 hpf (or heat shocked at 32 hpf) and analysed at 48 hpf. Scale bar: 20 μm. Rockout: Note that even at high Rockout concentration the microridges persist. The cell shape on the other hand is more irregular compared to the control. CK-666: Note the complete absence of microridges and reduced apical area of cells. NWASP-CA overexpression: This construct was injected mosaically and the affected cell is marked with arrowheads and is shown in the inset. Note the massively reduced apical area and altered structure of microridges. aPKC-CAAX overexpression: Note the altered shape of microridges and high accumulation of actin associated with adherens junctions compared to the control.

**Figure S4. Related to Figure 4 and 5.**
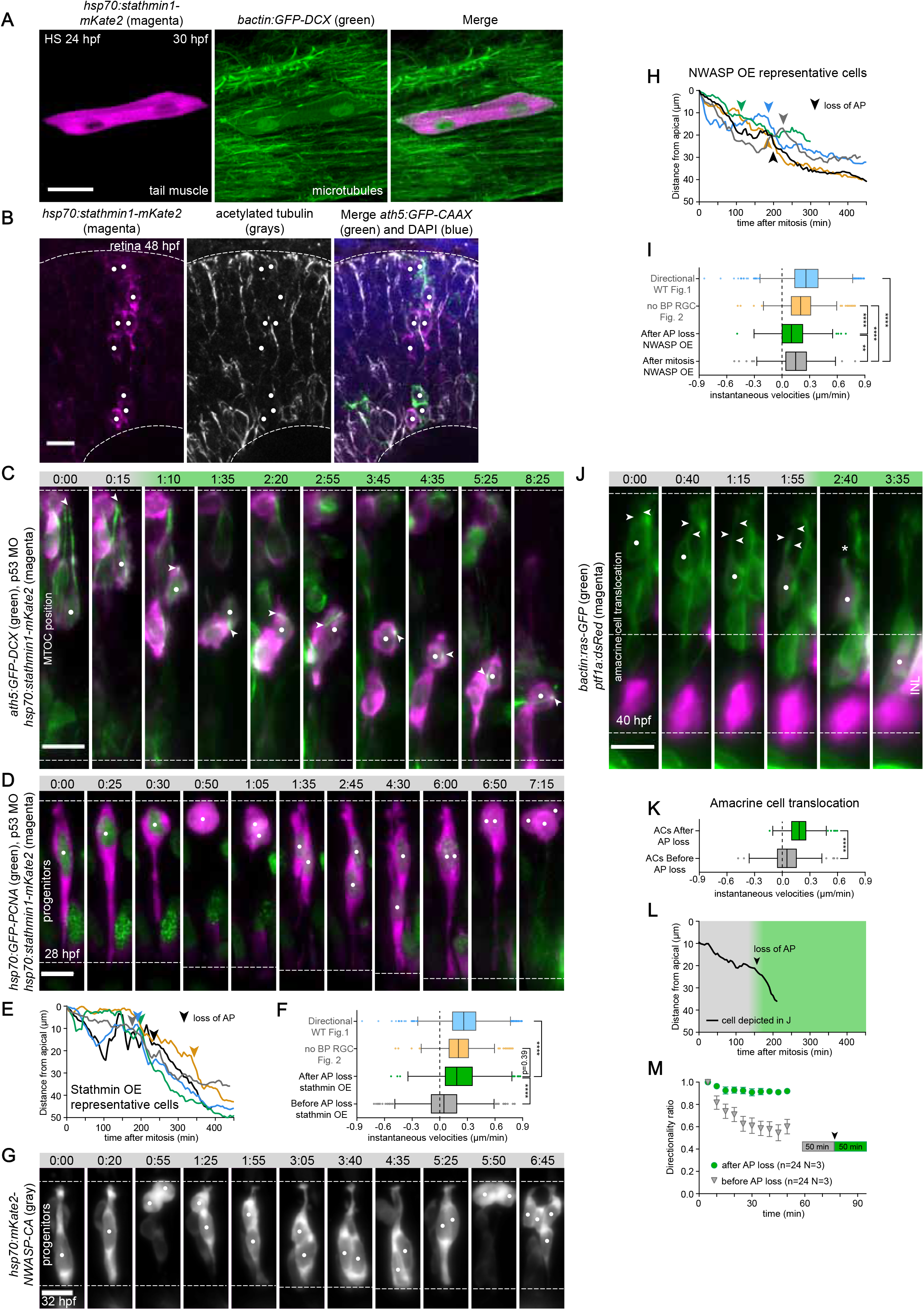
A) The destabilizing effect of the *hsp70:stathmin1-mKate2* construct on microtubules. The embryos of transgenic line ubiquitously expressing GFP tagged Doublecortin to label microtubules were injected with the *hsp70:stathmin1-mKate2*. The fish were heat shocked around 24 hpf and imaged live 6 hours later. Note the diffuse GFP-DCX signal and absence of microtubule filaments in the affected cell. Scale bar: 20 μm. B) Stathmin overexpression decreases microtubule acetylation. White dots: Stathmin overexpressing cells. Scale bar: 10 μm. C) Random microtubule organizing center (MTOC) position during RGC multipolar migration. Microtubules and MTOC were labeled by *ath5:GFP-DCX* injection. The RGC (white dot) initially has an apical MTOC (white arrowhead). The loss of apical process attachment triggers multipolar migratory mode (time 1:10) and later the MTOC is found at random position in the cell. Gray phase: cell still has the apical process. Green phase: directional multipolar mode. The dashed line delimits the apical and basal side of the retina. Time is shown in hh:mm. Scale bar: 10 μm. D) Microtubule depolymerization by *hsp70:stathmin1-mKate2* has no effect on retinal progenitor cell cycle or nuclear migration. The *ath5:gap-GFP* transgenic fish were injected with the *hsp70:stathmin1-mKate2* and *hsp70:GFP-PCNA* DNA to label nuclei. At 7:15 the *ath5:gap-GFP* expression starts. Time is shown in hh:mm. Scale bar: 10 μm. E) Five representative trajectories of the free migrating RGCs in the stathmin overexpression (OE) condition. 0 indicates the mitotic position of cells. The arrowhead marks the time point when the cell loses the apical process. F) Comparison of instantaneous velocities in stathmin overexpression before and during the multipolar migration and with control conditions. The instantaneous velocities are calculated from the one-dimensional movement along the apico-basal axis of the retina. The movement from apical to basal has a positive sign; the reverse movement has a negative sign. Directional WT Fig. 1 (n=140 N=24, 2587 data points), no BP RGC Fig. 2 (n=31 N=19, 448 data points), After AP loss stathmin OE (n=32 N=5, 631 data points), Before AP loss stathmin OE (n=32 N=5, 919 data points). Outliers were discarded by ROUT (Q=1.0%) for the plotting purposes, not for the statistical testing, which was done on the whole dataset. Statistical significance of the differences between instantaneous velocities: Directional WT Fig. 1 vs. After AP loss stathmin OE Mann-Whitney U test p<0.0001; After AP loss stathmin OE vs. Before AP loss stathmin Wilcoxon matched-pairs signed rank test p<0.0001; no BP RGC Fig. 2 vs. After AP loss stathmin OE Mann-Whitney U test p<0.3956. The data are shown as Tukey boxplot (box shows median and interquartile range and whiskers show 1.5 of the interquartile range). The median Directional WT Fig. 1=0.26 μm/min, no BP RGCs Fig. 2=0.20 μm/min, After AP loss stathmin=0.18 μm/min, Before AP loss stathmin=0.05 μm/min. G) Arp2/3 inhibition by *NWASP-CA* has no effect on retinal progenitor cell cycle or nuclear migration. WT fish were injected with the *hsp70:mKate2-NWASP-CA* and *ath5:GFP-CAAX* (not expressed in these cells) DNA. Time is shown in hh:mm. Scale bar: 10 μm. H) Five representative trajectories of the free migrating RGCs in the NWASP-CA overexpression (OE) condition. 0 indicates the mitotic position of cells. The arrowhead marks the time point when the cell loses its apical process. I) Comparison of instantaneous velocities in NWASP-CA overexpression before and during the multipolar migration and with control conditions. The instantaneous velocities are calculated from the trajectories of cells 95 min after mitosis and 95 after apical process loss. After AP loss NWASP OE (n=18 N=4, 342 data points), After mitosis NWASP OE (n=18 N=4, 342 data points). Outliers were discarded by ROUT (Q=1.0%) for the plotting purposes, not for the statistical testing, which was done on the whole dataset. Statistical significance of the differences between instantaneous velocities: Directional WT Fig. 1 vs. After mitosis NWASP OE Mann-Whitney U test p<0.0001; After AP loss NWASP OE vs. After mitosis NWASP OE Mann-Whitney U test p=0.0013; no BP RGC Fig. 2 vs. After AP loss NWASP OE Mann-Whitney U test p<0.0001; no BP RGC Fig. 2 vs. After mitosis NWASP OE Mann-Whitney U test p<0.0001. The data are shown as Tukey boxplot (box shows median and interquartile range and whiskers show 1.5 of the interquartile range). The median After AP loss NWASP OE=0.10 μm/min, After mitosis NWASP OE=0.15 μm/min. J) Amacrine cell (AC) translocation. *bactin:ras-GFP, ptf1a:dsRed* double transgenic embryos were used as donors for blastomere transplantation into WT acceptors. The translocating AC (white dot) initially has the AP (white arrowheads). Once it is lost (asterisk), this triggers fast directional movement to its final position. Gray phase: still with apical process. Green phase: directional multipolar mode. The dashed line delimits the apical side of the retina and the inner nuclear layer (INL), where the amacrine cells reside. Time is shown in hh:mm. Scale bar: 10 μm. K) Comparison of instantaneous velocities in amacrine cells (ACs) before and after apical process loss. 4 outliers were discarded by ROUT (Q=1.0%). ACs after AP loss (n=24 N=3, 257 data points), ACs before AP loss (n=24 N=3, 462 data points). The differences between instantaneous velocities are statistically significant: ACs after AP loss vs. ACs before AP loss Mann-Whitney U test p<0.0001. The median ACs after AP loss=0.18 μm/min, ACs before AP loss=0.05 μm/min. L) A representative trajectory of the translocating AC from (J). The arrowhead marks the time point when the cell loses the apical process. M) Directionality ratio of ACs before and after apical process loss. The average of all tracks is shown with error bars representing S.E.M. The interval of 50 min before and 50 min after AP loss was taken into account, because that is the average duration of the directional movement after apical process loss in ACs. Directionality ratio at the end of the trajectory: ACs after AP loss =0.92 ACs before AP loss =0.60.

**Figure S5. Related to Figure 6 and 7.**
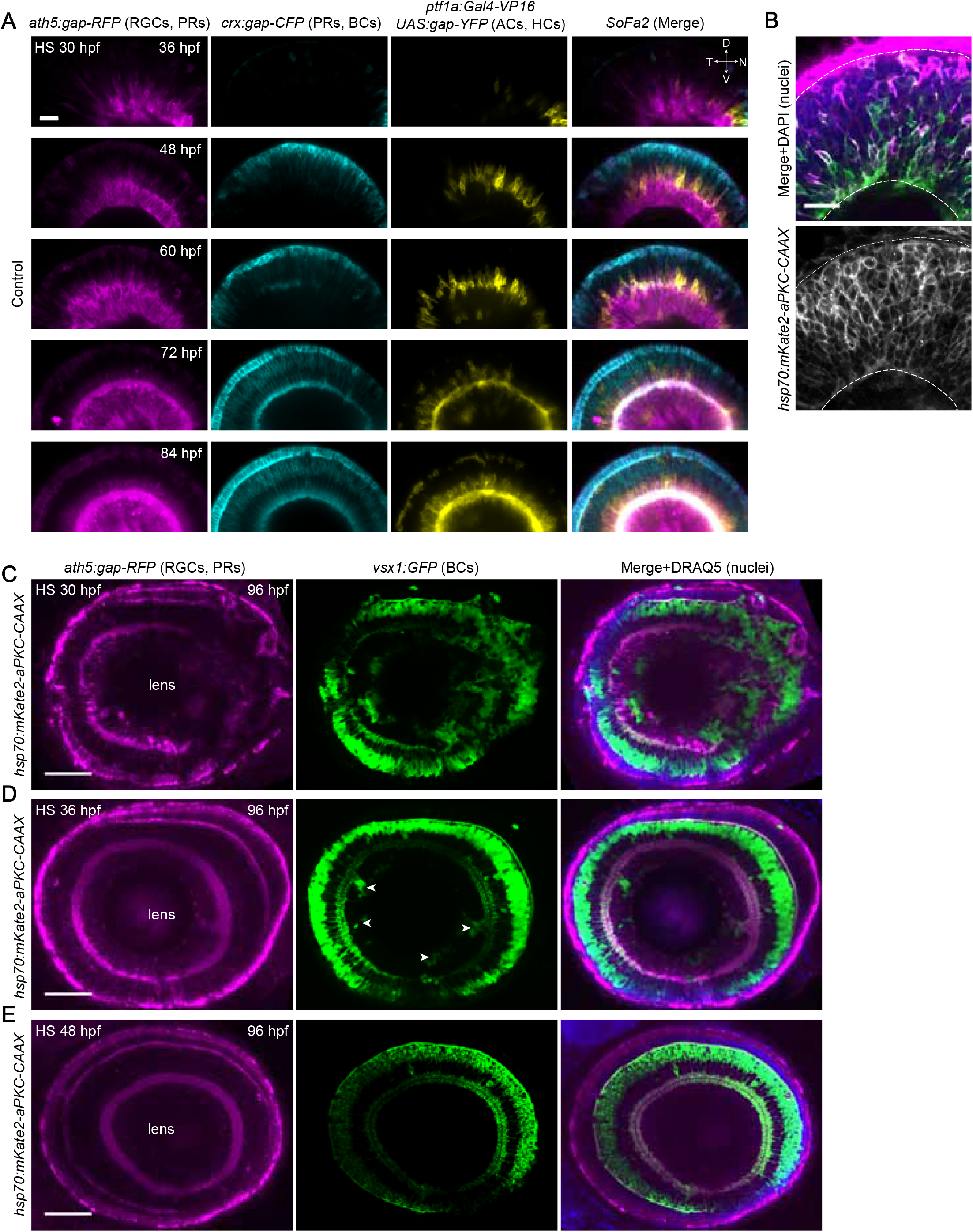
A) Control retina development. The *SoFa2* transgenic fish (combination of *ath5:gap-RFP* (labeling RGCs and photoreceptors), *crx:gap-CFP* (labeling photoreceptors and bipolar cells), *ptf1a:Gal4-VP16 UAS:gap-YFP* (labeling horizontal cells and amacrine cells) were heat shocked at 32 hpf and imaged in the LSFM from 36 hpf every 12 hours. The fish were kept in the incubator between the time points. No ectopic RGCs developed. Scale bar: 20 μm. B) Fluorescence signal of the *hsp70:mKate2-aPKC-CAAX* construct from Fig. 6 C. Fish were heat shocked at 30 hpf, fixed at 48 hpf and stained with Zn5 antibody. The fluorescence of mKate2-aPKC-CAAX is only detectable within 24 hours of heat shock due to the fast turnover of the protein. The dashed lines mark the apical and basal membranes of the retina. Scale bar: 20 μm. C) D) E) aPKC-CAAX perturbs retinal lamination only during a short time window. *ath5:gap-RFP, vsx1:GFP, hsp70:mKate2-aPKC-CAAX* triple transgenic line was heat shocked at 30 hpf (C), 36 hpf (D) or 48 hpf (E), fixed at 96 hpf and stained with DRAQ5 to additionally label the nuclei. The severe lamination problems can be observed in (C). Mild phenotype can be seen in (D) with only a few ectopic clusters of bipolar cells (arrowheads) and in (E) the retina is indistinguishable from the control (Fig. 7 A) with only one continuous RGC layer and the whole retina forming an unperturbed layered structure. Scale bar: 50 μm.

